# The ABC-F protein Uup, Rescues Stalled Ribosomal Termination Complexes to Modulate DNA Maintenance

**DOI:** 10.64898/2026.10.08.757592

**Authors:** Agata Wawszczyk, Saravuth Ngo, Marion Hamon, Laura Monlezun, Grégory Boël

## Abstract

The cytosolic ATP-binding cassette F (ABC-F) family modulates ribosomal activity in eukaryotes and prokaryotes. Although they all bind to the E site of the ribosome, some bacterial ABC-Fs confer resistance to ribosome-targeting antibiotics, while housekeeping members have poorly defined roles. Interestingly, in *Escherichia coli,* which possesses four ABC-F (EttA, Uup, YbiT and YheS), Uup has been rather implicated in DNA recombination. Using an unbiased approach, we reconcile this apparent discrepancy by showing that Uup’s involvement in DNA recombination is indirect and is dictated by Uup-mediated regulation of the last steps of translation that rescues problematic translation of genes involved in DNA maintenance such as *ssb* and *dnaX*. Accordingly, the elongated cell phenotype observed in *uup* deletion background is reverted by expressing a *ssb* variant that no longer depends on Uup. Unlike its paralog EttA, which helps ribosomes in translating proteins with repeats of negatively charged residues in the N-terminal 30 amino acids, Uup instead facilitates the rescue of stalled translation termination events that, in its absence, are handled by the tmRNA pathway.

## Introduction

The *uup* gene was first characterized in a genetic screen in *E. coli* aimed at identifying mutations that increased the frequency of precise excision of Tn5 and Tn10 transposons, as well as Insertion Sequences (IS) elements (Hopkins *et al*, 1983). A subsequent screen, focused specifically on Tn10 precise excision, further characterized mutations in *uup* and identified additional mutations in the *ssb* gene (Reddy & Gowrishankar, 1997). Precise excision of transposons is a host-mediated deletion event that occurs at short direct repeats generated during transposition by the large inverted repeats that constitute the IS of these elements (Foster *et al*, 1981). This process is independent of both the host *recA* function and transposon-encoded functions (Foster *et al*., 1981; Reddy & Gowrishankar, 1997). These screens demonstrated that mutations in *uup* or *ssb* increase the frequency of precise transposon excision. The *ssb* gene encodes the single-stranded binding protein (SSB) (Meyer *et al*, 1979), which is essential for DNA replication, recombination, and repair (Chase & Williams, 1986; Meyer & Laine, 1990). SSB binds cooperatively as a tetramer to single-stranded DNA, protecting it during replication and recruiting partner proteins via its C-terminal extension to sites of DNA metabolism (Shereda *et al*, 2008).

The phenotype of *ssb* mutants regarding transposon release can be explained by SSB’s role in DNA repair and recombination. SSB binds to single-stranded DNA (ssDNA), and protects it from nuclease activity by sequestering it, thereby reducing its reactivity (Molineux & Gefter, 1975; Myler *et al*, 2016). For *uup* mutants, the explanation remains more elusive. Uup belongs to the ABC-F protein family (part of the ATP Binding Cassette superfamily), characterized by two tandem ABC ATPase domains connected by a linker that contains the P-site tRNA interaction motif (Boel *et al*, 2014) (PtIM; PFAM domain PF12848). ABC-F proteins share this ATPase domain architecture with other ABC proteins, including some ABC transporters as well as the UVR family; however, the PtIM motif is exclusive to the ABC-F protein family (Ousalem *et al*, 2019). Since most members of the UVR family (e.g., RecA, Rad50) are involved in DNA repair processes, their structural homology with Uup suggests that Uup may also function in DNA repair, potentially explaining the phenotype of transposon release through direct action on DNA (Murat *et al*, 2006; Romero *et al*, 2020a; Romero *et al*, 2020b).

Dassa and Cox laboratories have conducted studies to demonstrate a direct action of Uup on DNA (Burgos Zepeda *et al*, 2010; Carlier *et al*, 2012; Murat *et al*., 2006; Romero *et al*., 2020a; Romero *et al*., 2020b). The Dassa group first showed that an ATPase-deficient mutant of Uup increases the release of the Tn10 transposon (Murat *et al*., 2006), suggesting that the ATPase activity is essential for Uup function. Secondly, they showed that Uup binds to double-stranded DNA *in vitro*, and that a deletion of its ∼80-residue C-terminal extension, which belongs to the PF16326 domain family, reduces its affinity for DNA (Murat *et al*., 2006). Additionally, the C-terminal extension alone was found to bind DNA and to be essential for the phenotype of transposon release (Burgos Zepeda *et al*., 2010). This C-terminal extension forms a coiled-coil motif, as determined by NMR (Carlier *et al*., 2012).

The Cox group, used an elegant plasmid-based deletion assay where the tetracycline gene is inactivated by a repeat of its sequence, to demonstrate that the deletion of *uup* (Δ*uup*) or *radD*, a RecQ-like SF2 family ATPase, increases the generation of functional tetracycline recombination events (Romero *et al*., 2020a). Both deletions also increase cell size and Δ*uup* cells showed DNA partitioning defects (Romero *et al*., 2020a). Surprisingly, the study showed that Uup binds to Holliday junction structures *in vitro,* at high concentrations exclusively, but not to dsDNA, contrasting earlier findings from the Dassa laboratory (Burgos Zepeda *et al*., 2010; Murat *et al*., 2006). They also revealed an epistatic relation between deletions of *uup* and *recG*, a DNA helicase involved in double-strand break repair.

On the other hand, ABC-F proteins are well known for their interaction with the ribosome by binding to its E-site, as demonstrated by several structural studies of ABC-F proteins in complex with ribosomes from diverse organisms (Boel *et al*., 2014; Chen *et al*, 2014; Crowe-McAuliffe *et al*, 2018; Crowe-McAuliffe *et al*, 2022; Crowe-McAuliffe *et al*, 2021; Mohamad *et al*, 2022; Obana *et al*, 2023; Singh *et al*, 2023; Su *et al*, 2019; Su *et al*, 2018). As translation factors, ABC-F proteins modulate ribosome activity during mRNA translation facilitating the translation of mRNAs encoding particular amino acid motifs challenging for the ribosome to synthesize (Boel *et al*., 2014; Chadani *et al*, 2024; Hong *et al*, 2024; Ousalem *et al*, 2024; Ousalem *et al*, 2023; Singh *et al*., 2023; Takada *et al*, 2024). Indeed, long considered passive bystanders during translation, nascent polypeptides have recently been shown to stall the ribosome depending on their amino acid sequences. This phenomenon, defined as Context-Dependent Ribosomal Stalling (CDRS), is now recognized as a significant factor in the regulation of protein synthesis, and some ABC-F proteins act as modulators of specific CDRS events.

We demonstrated that one of the four ABC-F protein in *E. coli* named EttA, increases the translation of specific mRNAs. Some of these mRNAs encode enzymes of the Krebs cycle’s glyoxylate shunt pathway. Furthermore, identification of specific EttA targets has revealed a peptide signature, namely, the presence of negatively charged residues in the early coding sequence (Ousalem *et al*., 2024). Another study using non-physiological genetic reporters has reached the same conclusion (Chadani *et al*., 2024).

Concerning the three other ABC-F of *E. coli* (Uup, YbiT and YheS), their physiological function in mRNA translation process is much less understood, though each factor is proposed to have a unique cellular function (Chadani *et al*., 2024; Ousalem *et al*., 2024; Ousalem *et al*., 2023; Singh *et al*., 2023). Primarily based on *in vitro* experiments, it has been proposed that YheS may rescue ribosomes stalled at the SecM stalling sequence, while YbiT may act on sequences containing positively charged residues (Chadani *et al*., 2024). Regarding Uup, it has been shown to increase the expression of a genetic reporter with a 10-residue polyproline segment at its C-terminal end *in vitro*. Morever, addition of Uup reduces peptidyl-tRNA formation, which arises from the abortive translation of the reporter (Chadani *et al*., 2024). Uup can also partially complement a ribosome biogenesis defect caused by deletion of the *bipA* gene (Cochrane, 2015; Murina *et al*, 2019). Additionally, under certain conditions, Uup is necessary for optimal cellular fitness (Murat *et al*, 2008; Ousalem *et al*., 2023); a requirement also observed for EttA (Boel *et al*., 2014). In *Bacillus subtilis*, it has been shown that YfmR, which seems to be a Uup orthologue, can rescue ribosome stalling on polyproline stretches (Hong *et al*., 2024), whereas YkpA, a YbiT homologue, promotes the translation of positively and negatively charged motifs (Takada *et al*., 2024).

One key tool used to determine the structures of ABC-F proteins in complex with the ribosome, relies on ABC-F mutants carrying dual glutamate-to-glutamine mutations (EQ_2_) in the catalytic residues following the Walker B motifs in both ABC domains (Boel *et al*., 2014; Chen *et al*., 2014). These mutations prevent ATP hydrolysis, effectively trapping the ABC-F proteins in a closed, ATP-bound conformation on the ribosome (Boel *et al*., 2014; Chen *et al*., 2014; Crowe-McAuliffe *et al*., 2018; Crowe-McAuliffe *et al*., 2022; Crowe-McAuliffe *et al*., 2021; Mohamad *et al*., 2022; Obana *et al*., 2023; Singh *et al*., 2023). Although all EQ_2_ mutants inhibit protein synthesis (Murina *et al*., 2019; Ousalem *et al*., 2023), they act at different stages of translation: EttA and YbiT inhibit the first steps of elongation, while YheS and Uup inhibit later steps (Ousalem *et al*., 2023).

Finally, a class of specialized ABC-F, termed ARE-ABC-F (<u>a</u>ntibiotic <u>re</u>sistance), confers resistance to antibiotics that bind to the large (50S) subunit of the ribosome (Fostier *et al*, 2021). These proteins bind to the ribosome in a manner similar to other ABC-F proteins (Crowe-McAuliffe *et al*., 2018; Crowe-McAuliffe *et al*., 2022; Crowe-McAuliffe *et al*., 2021; Mohamad *et al*., 2022; Obana *et al*., 2023; Su *et al*., 2018) and are capable of alleviating ribosomes stalled by antibiotics (Fostier *et al*., 2021; Fostier *et al*, 2023; Lenart *et al*, 2015; Sharkey *et al*, 2016; Vimberg *et al*, 2020). Additionally, they can regulate their own expression, as well as that of other genes involved in antibiotic synthesis, by rescuing ribosome pausing on regulatory leader peptides (Fostier *et al*., 2023; Koběrská *et al*, 2025; Koberska *et al*, 2021).

To reconcile the apparent discrepancy between the functions of Uup in DNA maintenance and mRNA translation, we employed a comprehensive, unbiased approach using proteomic and transcriptomic analyses following the characterization of the phenotypes of the *uup*-deleted strain. We identified changes in gene expression in the Δ*uup* strain and validated several targets using fluorescent genetic reporters. Two of the validated targets, *ssb* and *dnaX*, are involved in DNA maintenance processes. We demonstrated that the changes in expression result from impaired translation of their mRNA, which occurs after stalling of the ribosome at the termination step. Stalled terminating ribosomes trigger the *ssrA* response, tagging the ribosomal elongating peptide for degradation by the cellular degradation machinery. The decreased expression of SSB in the Δ*uup* strain appears to explain the long-cell phenotype induced by DNA instability observed for Δ*uup* strain. In conclusion, our work suggests that the primary effect of Uup is on mRNA translation, and the reduced expression of specific proteins in the absence of Uup leads to the DNA maintenance phenotypes previously observed in the Δ*uup* strain.

## Results

### uup is required for genome stability and resistance to UV-induced damage

Previous results (Romero *et al*., 2020a) have shown that *uup* deletion alters cell morphology by causing incomplete nucleoid separation. We first characterized the cell shape of our Δ*uup* strain by light microscopy (**Fig. 1A** and **EV Fig 1A**). The Δ*uup* strain in late exponential phase showed heterogeneous cell lengths. Quantification indicates that the average length is 2.8 µm for the WT strain and 3.5 µm for the Δ*uup* strain (**Fig. 1B**); some Δ*uup* cells reached up to 15 µm in length. DAPI staining revealed that in elongated Δ*uup* cells the chromosome is less condensed than in WT cells, suggesting incomplete nucleoid separation.

**Figure 1:**
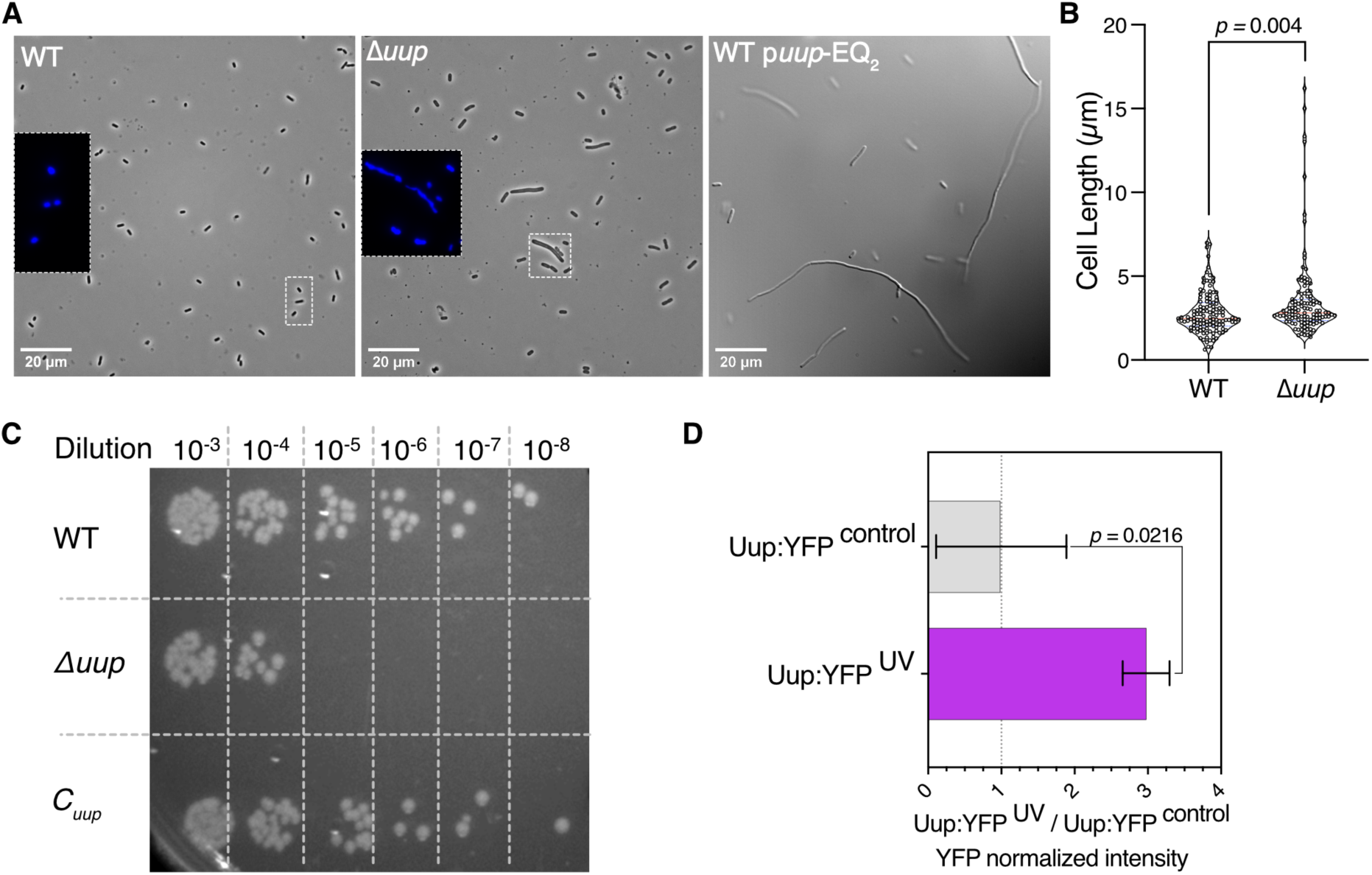
Deletion of *uup* induces phenotypes related to genome instability. **A)** Phase-contrast microscopy with DAPI staining of the WT (left) and Δ*uup* (middle) strains from LB culture at OD_600_ = 0.7. Bright-field microscopy of the strain expressing the ATPase-deficient Uup-EQ_2_ variant on a plasmid, induced with 0.1% L-Arabinose, collected after growth inhibition caused by Uup-EQ_2_ (right). **B**) Violin plot showing cell length measurements for the WT (n = 123) and Δ*uup* (n = 116) strains. Measurements were performed using MicrobeJ (Ducret *et al*., 2016). **C**) Drop dilution assay of WT, Δ*uup*, and C*_uup_* strains after UV exposure. **D**) Fluorescence of the *uup:yfp* genomic fusion in cultures at OD_600_ = 1, 5.5 h after UV exposure (4 500 000 μJ / cm^2^ UV at 254 nm). Histograms show fluorescence normalized to non-exposed cells (Uup:YFP^control^). B-D) Error bars represent the mean ± s.d. for triplicate experiments. *P*-values were calculated using an unpaired two-tailed *t*-test.

Interestingly, expression from a plasmid of the ATPase-deficient Uup mutant (Uup-EQ_2_) caused, as previously reported (Murina *et al*., 2019; Ousalem *et al*., 2023), inhibition of cell growth. However, when we observed cells by microscopy just after growth inhibition (**Fig. 1A, right**), they were extremely elongated, reaching lengths greater than 80 µm. Thus, the elongated-cell phenotype of the Δ*uup* strain can be further exacerbated by expressing Uup-EQ_2_ in the WT strain. This phenotype was previously observed and described as the extremely elongated (eel) cells phenotype (El-Hajj Ziad & Newman Elaine, 2015), attributed to low expression of the cell division protein FtsZ; it was also reported after UV irradiation (Long *et al*, 1978) and following nalidixic acid treatment (Goss William *et al*, 1964). This suggests that DNA damage can induce the long-cell phenotype, reinforcing the idea that Uup is involved in DNA maintenance.

Prompted by these observations, we tested the survival of the WT and Δ*uup* strains after UV exposure, as well as that of the complemented strain, carrying the WT *uup* gene inserted at the P21 locus under the control of its endogenous promoter (C*_uup_*). The Δ*uup* strain shows increased sensitivity to UV_254 nm_ (**Fig. 1C**). After UV exposure, no colonies formed from the 10^−5^ dilution of the Δ*uup* culture, whereas colonies were observed up to the 10^−8^ dilution for the WT and complemented strains. Pre-exposure controls (**EV Fig. 1B**) showed equivalent colony formation for the different strains, although the Δ*uup* strain produced slightly fewer colonies. This is likely because Δ*uup* cells are longer and therefore there are fewer cells per OD_600_ unit. We also assessed Uup expression in a strain expressing Uup fused to yellow fluorescent protein (*uup*:*yfp*). The YFP fluorescence of the UV-treated culture was higher than that of the non-treated culture (**Fig. 1D**). No difference in fluorescence was observed between the cultures prior to UV exposure (**EV Fig. 1C**). These results demonstrate that Uup is important for UV protection and that its expression is induced by UV.

### ATPase-deficient Uup (Uup-EQ_2_) induces ribosome stalling during late elongation or termination

The Uup paralogue EttA acts early during translation (Chadani *et al*., 2024; Ousalem *et al*., 2024) accordingly, its ATPase-deficient mutant EttA-EQ_2_ blocks ribosomes at initiation or during the earliest stages of elongation (Boel *et al*., 2014; Chen *et al*., 2014; Ousalem *et al*., 2024). A sucrose-gradient profile of cell extracts from a strain expressing His_6_-EttA-EQ_2_ in exponential phase showed a decrease in polysomes and an increase in 70S and disome peaks (**Fig. 2**). Ribosome-associated His_6_-EttA-EQ_2_ were enriched in the 70S and disome fractions in this condition. By contrast, expression of His_6_-Uup-EQ_2_ produced the opposite profile: increased polysomes, decreased 70S, and most ribosome-associated His_6_-Uup-EQ_2_ found in polysome fractions (**Fig. 2**). These results demonstrate that Uup-EQ_2_ primarily blocks ribosomes during late elongation or termination: stalled ribosomes lead to accumulation of trailing ribosomes, resulting in increased polysome formation.

**Figure 2:**
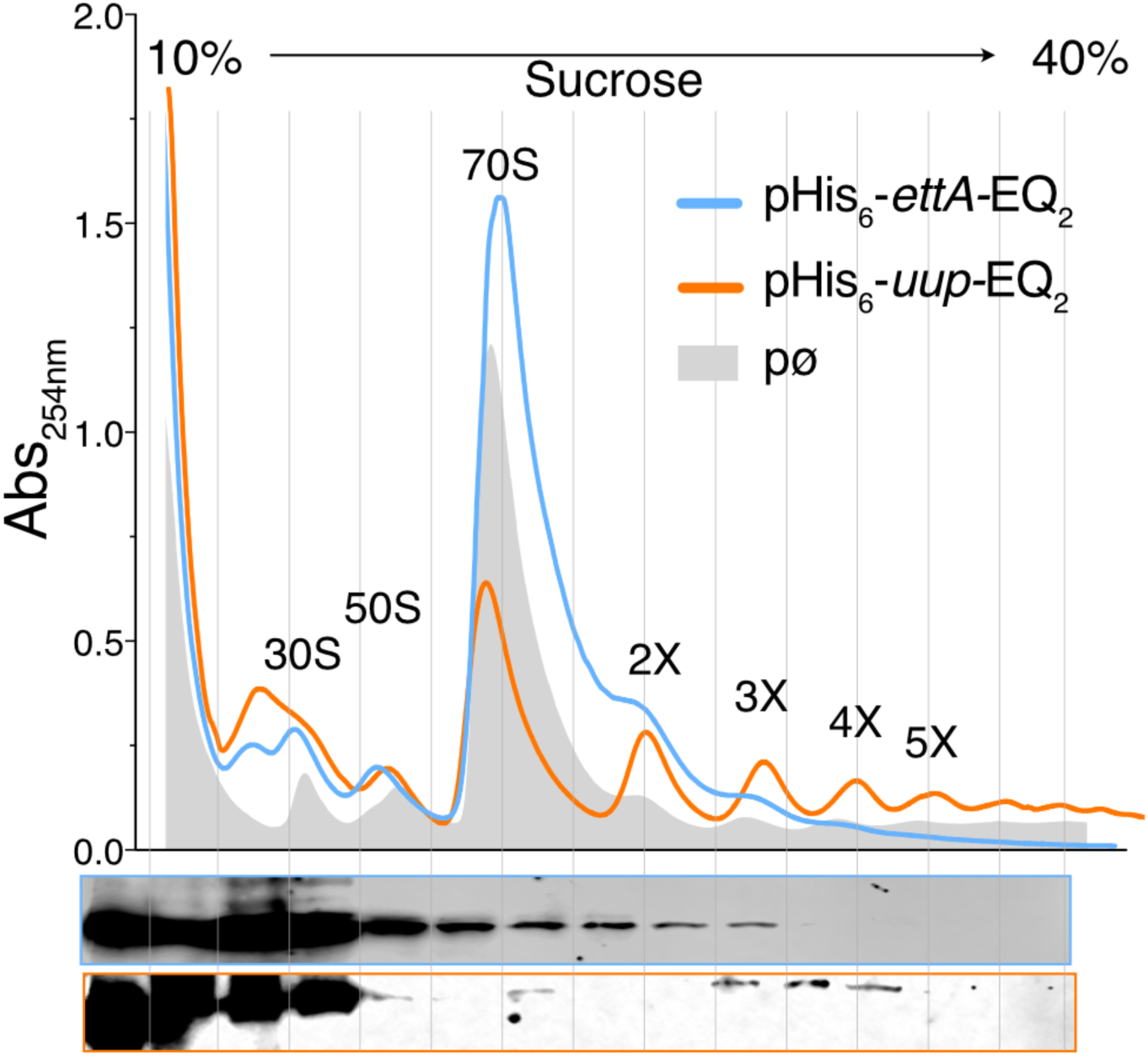
ATPase-deficient Uup-EQ_2_ induces polysome accumulation, while EttA-EQ_2_ induces depletion. **A**) Polysome profiling on a 10–40% sucrose gradient of lysates from exponential-phase cultures of WT strains expressing His_6_-Uup-EQ_2_ or His_6_-EttA-EQ_2_ on a pBAD plasmid induced with 0.1% L-Arabinose. The control polysome profile of a strain transformed with an empty pBAD plasmid is shown as a grey shadow. Below, western blot analysis of the fractions probed with an anti-His antibody, the first lane is the input.

### Proteomic and transcriptomic analysis show reduce expression of proteins involved in DNA replication and induction of the SOS response

Based on these results and previous ones (Ousalem *et al*., 2023; Singh *et al*., 2023), we hypothesized that Uup may be involved in the translation of specific mRNAs. To identify translation events modulated by Uup, we conducted proteomic (**EV Fig. 2**) and transcriptomic analyses of the WT and Δ*uup* strains. Analysis of both datasets (**Fig. 3**) revealed an increase in the SOS response (Matic *et al*, 1995; Sass & Lovett, 2024) in the Δ*uup* strain. While some upregulation is observed at the protein level, the effect is more pronounced in the transcriptomic data, likely because transcripts are detected more comprehensively than proteins. Interestingly, some genes involved in the SOS response, such as *dnaQ*, show greater induction at the protein level than at the RNA level. The SOS response is a global transcriptional program triggered by DNA damage, leading to the expression of DNA repair enzymes (Matic *et al*., 1995; Sass & Lovett, 2024). This observation confirms that deletion of *uup* causes major DNA integrity stress in the cell. In addition to this regulon, several pH-inducible genes, such as *gadA* and *gadB* (glutamate decarboxylases A and B), the related L-glutamate:4-aminobutyrate antiporter *gadC*, and the starvation lipoprotein *slp* are also upregulated. Inversely, deletion of *uup* reduces the expression of key proteins involved in DNA maintenance and replication, such as the single-stranded DNA-binding protein (SSB) and the DNA polymerase III γ subunit (DnaX), as well as other proteins like the lipoprotein NlpA. This reduction is more pronounced at the protein level than at the RNA level.

**Figure 3:**
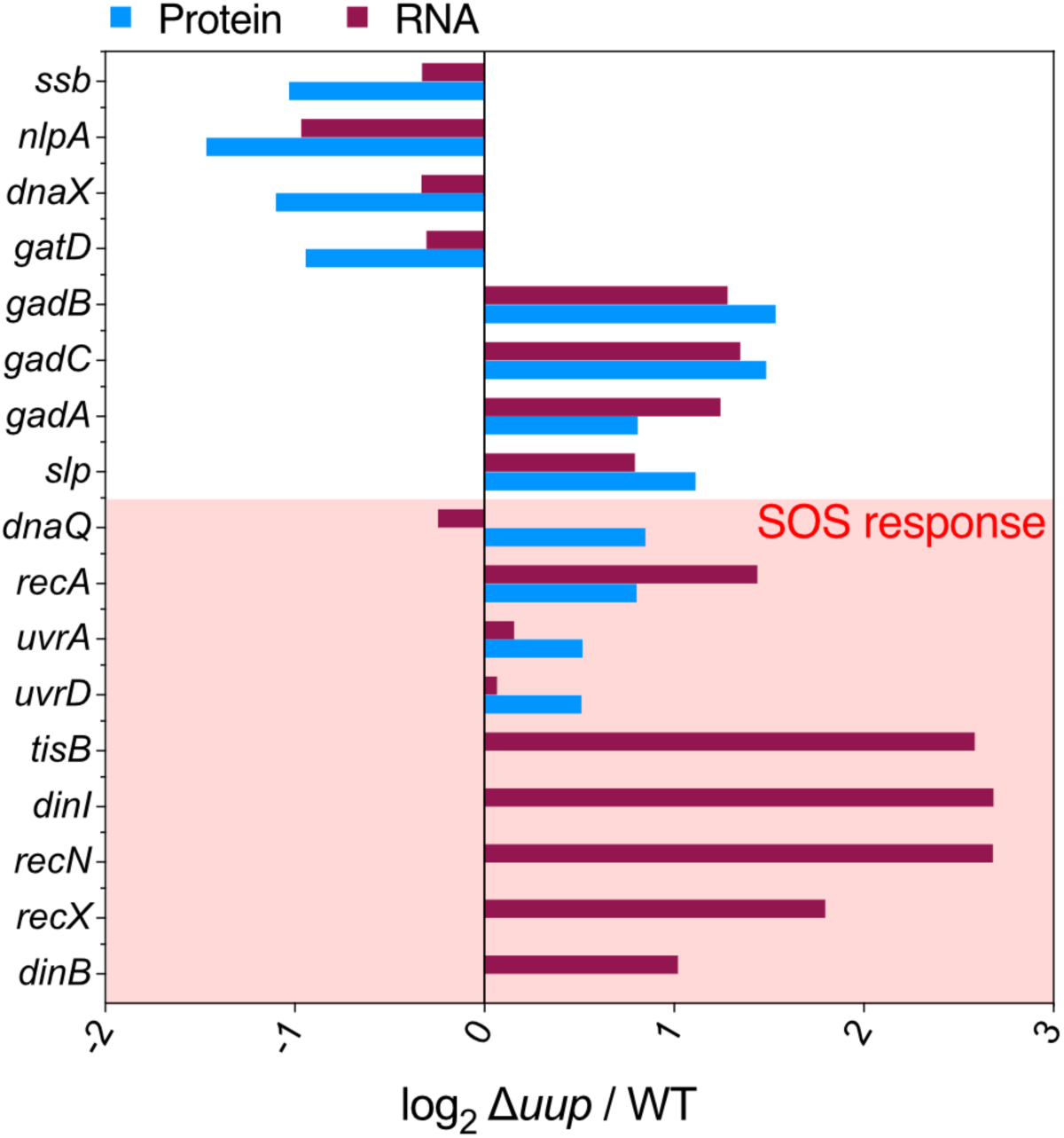
**Deletion of the *uup* gene represses the expression of SSB, NlpA, and DnaX proteins while inducing an overall transcriptional increase of SOS response genes.** Histograms showing the log_2_ Δ*uup* / WT ratio of selected proteins or mRNAs identified in proteomic and transcriptomic studies conducted on cell cultures collected at OD_600_ = 2. For some of them, quantification can be determined for the mRNA only.

### uup-dependent regulation involves the C-terminal residue of target proteins

To confirm the translational regulation of some of the genes downregulated in the Δ*uup* strain, we used YFP translational reporters. These reporters, containing either part or the full sequence of the tested gene, were expressed from the low-copy plasmid pMMB67EH (Fürste *et al*, 1986) under the control of the IPTG-inducible pLlacO promoter. Each reporter was transformed into the WT, Δ*uup*, and C*_uup_* strains.

We first constructed *ssb* reporters by fusing the region encoding its last 50 amino acids (aa) either upstream or downstream of the *yfp* gene. For the upstream fusion, a methionine start codon was added at the beginning of the *ssb* fragment. The upstream *ssb* reporter (*ssb*_129-178aa_:*yfp*) showed low YFP fluorescence, with no difference in signal between the three strains (**Fig. 4A and EV Fig. 3A**). In contrast, the downstream fusion (*yfp*:*ssb*_129-178aa_) exhibited a higher fluorescence signal (**EV Fig. 3A**), which was specifically reduced in the Δ*uup* strain (**Fig. 4A**). This confirms that Uup affects the expression of this reporter, likely at the translational level, and that this effect occurs only when translation terminates with the *ssb* sequence.

**Figure 4:**
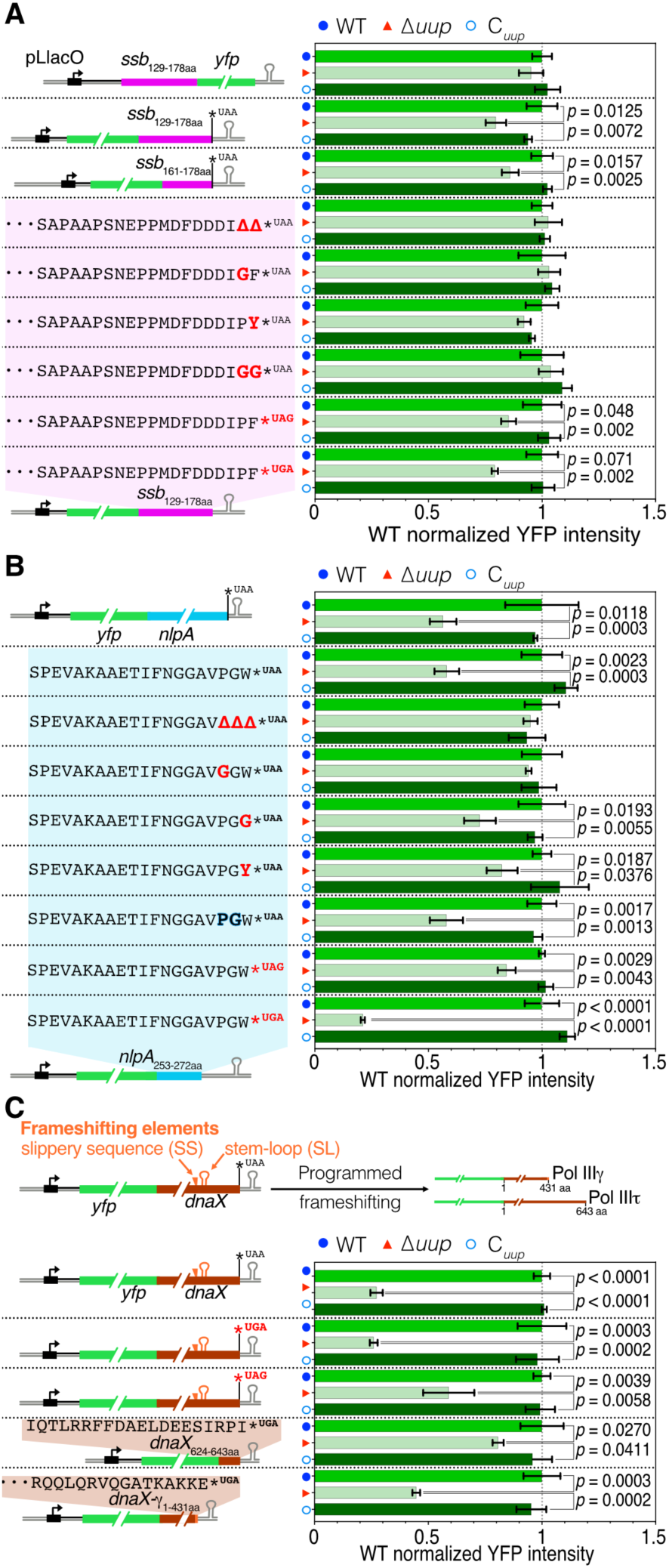
Uup facilitates translation of *ssb*, *nlpA*, and *dnaX* mRNAs, this regulation depends on the C terminal residue encoded by the mRNA and on the identity of the stop codon. Histograms show fluorescence ratios normalized to the wild type (WT) strain for YFP translation reporters expressed from a low copy plasmid in WT, Δ*uup*, and complemented (C*_uup_*) strains after 30 hours of growth in microplate (full growth curves and fluorescences are presented in **EV** fig. 3). Error bars represent mean ± s.d. from triplicate experiments, calculated as described in the methods section. *P*-values were obtained using an unpaired two-tailed *t*-test. **A)** YFP reporters containing portions of the *ssb* coding sequence. Constructs include the last 50 codons (*ssb*_129-178aa_) placed either upstream or downstream of YFP, the last 18 codons (*ssb*_161-178aa_) placed downstream of YFP, and variants of the 50-codon construct (*ssb*_129-178aa_) placed downstream of YFP in which the terminal residues are mutated and the stop codon is replaced by one exclusively recognized by RF1 (UAG) or RF2 (UGA). **B**) YFP reporters containing either the full *nlpA* sequence or the last 20 codons (*nlpA*_253-272aa_) placed downstream of YFP. For the 20-codon construct, variants in which the terminal residues are mutated (in bold and red) or replaced by synonymous codon (in bold and blue) or in which the stop codon is replaced by one exclusively recognized by RF1 (UAG) or RF2 (UGA) are depicted. **C**) YFP reporters containing the full *dnaX* sequence encoding the tau (Pol IIIτ) and gamma (Pol IIIγ) subunits of DNA polymerase III. The γ subunit is produced by a −1 Programmed Ribosomal Frameshift (−1PRF) that occurs at a poly(A) tract forming a slippery sequence (SS) and generates a new stop codon immediately following the frameshift signal. The −1PRF also relies on a stem-loop (SL) downstream of the SS. Variants in which the non-frameshifted stop codon was replaced by one recognized exclusively by RF1 (UAG) or RF2 (UGA) were tested. Variant with the last 20 codons of Pol IIIτ (*dnaX*_624-643aa_) was also tested. Finally, construct that produces only the γ subunit was generated by inserting an additional adenine into the slippery sequence and truncating the *dnaX* sequence after the stop codon read in the frameshifted reading frame (*dnaX*-γ_1-431aa_). The full growth curves with fluorescence intensity are presented in **EV Fig. 3**.

We then reduced the *ssb* sequence to the last 18 aa of the coding sequence (*yfp*:*ssb*_161-178aa_) and observed that the specific downregulation in the Δ*uup* strain persisted, though it was less pronounced. Deletion of proline (Pro) and phenylalanine (Phe), the last two residues of SSB, in the *yfp*:*ssb*_129-178aa_ construct, abolished Uup-dependent expression, as did the replacement of Pro with glycine (Gly) or of Phe with tyrosine (Tyr). As expected, simultaneously replacing the last two residues with two Gly residues also abolished Uup-dependent expression.

Additionally, we noticed that the nature of the stop codon could influence the amplitude of the response. Indeed, the native stop codon of *ssb* is UGA, whereas our original constructs used UAA. *E. coli* can exploit two specific release factors (RFs), RF1 and RF2, which bind to the ribosomal A site when a stop codon is present and hydrolyze the ester bond between the P-site tRNA and the synthesized protein, thereby releasing it. While both RF1 and RF2 can recognize the UAA stop codon, UAG is specifically decoded by RF1, and UGA by RF2. When we replaced the UAA stop codon with UAG, the decrease in expression in the Δ*uup* strain, compared to the WT or C*_uup_* strains, was similar to that observed with the UAA codon (**Fig. 4A and EV Fig. 3A**). However, replacing it with the RF2-dependent UGA codon increased the amplitude of the decrease (**Fig. 4A and EV Fig. 3A**). These observations suggest that Uup-dependent expression of *ssb* is more pronounced when translation termination involves an RF2-dependent stop codon.

We applied the same approach to *nlpA* (**Fig. 4B and EV Fig. 3B**) and observed similar results. First, combining the *yfp* gene with the full *nlpA* sequence in place of its stop codon resulted in reduced expression specifically in the Δ*uup* strain. Reducing the *nlpA* sequence to the last 20 aa (*yfp:nlpA_253-272aa_*) (with the addition of a methionine as the first codon) showed a similar reduction in expression in the Δ*uup* strain. Deletion of the last three residues (PGW) abolished the differential expression in the Δ*uup* strain, as did the replacement of the Pro with Gly. Replacing the terminal Trp with Gly or Tyr reduced the amplitude of the differential expression (**Fig. 4B and EV Fig. 3B**). These results highlight the importance of the last amino acids in the expression change, particularly the terminal Pro.

Moreover, replacing the native Pro and Gly codons (CCU and GGC respectively) of the terminal PGW motif with synonymous codons (CCG and GGG respectively) did not affect the regulation, suggesting that the regulation is driven more by an amino acid effect than an RNA effect (**Fig. 4B and EV Fig. 3B**). When the native UAA stop codon of *nlpA* was changed to the RF1-dependent UAG codon, the effect on Uup-dependent expression was reduced by 1.5-fold whereas changing it to the RF2-dependent UGA codon increased the dependency on Uup by more than twofold (2.8-fold).

Finally, we tested the expression of the *dnaX* gene, which encodes two distinct proteins: the DNA polymerase III γ subunit (Pol IIIγ) and the DNA polymerase III τ subunit (Pol IIIτ). This dual expression is achieved through −1 programmed ribosomal frameshifting (−1PRF) (Blinkowa & Walker, 1990; Korniy *et al*, 2019). A slippery sequence (SS), composed of the heptanucleotide A AAA AAC, along with a downstream RNA stem-loop (SL), can induce the ribosome to shift its reading frame by −1 nucleotide (**Fig. 4C**) (Caliskan *et al*, 2017; Kim *et al*, 2014; Larsen *et al*, 1997). Without frameshifting, the ribosome synthesizes the 643 aa Pol IIIτ protein. However, when frameshifting occurs at the SS, the ribosome shifts backward by one nucleotide (−1 frame), resulting in the read of an extra adenosine in the SS. This frameshift leads to the synthesis of the 431 aa Pol IIIγ protein, which shares the same sequence as Pol IIIτ but is truncated by the last 213 aa, with its terminal amino acid changing from serine (Ser) to glutamate (Glu).

We first tested the expression of the full *dnaX* sequence fused downstream of the *yfp* gene (which leads to the production of two proteins: YFP-Pol IIIγ and YFP-Pol IIIτ) in the three strains (**Fig. 4C and EV Fig. 3C**). Expression in the Δ*uup* strain was reduced by 3.7-fold compared to the WT and C*_uup_* strains. Our initial construct used a UAA stop codon, which is not the native stop codon of *dnaX*. We therefore tested an equivalent construct with a UGA stop codon, matching the native sequence of *dnaX*. This construct showed the same decrease in expression in the Δ*uup* strain. However, when we replaced the RF2-dependent UGA stop codon with the RF1-dependent UAG stop codon, we observed only a 1.2-fold decrease in expression in the Δ*uup* strain. This result, also observed for *nlpA* and *ssb*, confirms the importance of the stop codon in Uup-dependent expression, with RF2-decoded stop codons amplifying the differential expression.

To determine whether the effect of Uup was specific to the synthesis of Pol IIIτ or could also be true for Pol IIIγ through −1PRF, we fused either the sequence encoding the last 20 aa of Pol IIIτ (*yfp*:*dnaX*_624-643aa_) or the sequence (*yfp*:*dnaX*-γ_1-431aa_) encoding Pol IIIγ (with an additional adenosine in the SS of the frameshifting element to mimic the translation during −1PRF) to the end of the *yfp* gene. Importantly, this Pol IIIγ construct lacked the SL necessary for −1PRF to occur. The reporter containing the sequence encoding the last 20 aa of Pol IIIτ showed a 1.2-fold decrease in expression in the Δ*uup* strain compared to the WT and *C_uup_* strains, while the reporter containing the sequence encoding Pol IIIγ showed a 2.2-fold decrease. These results demonstrate that Uup-dependent expression occurs for both reporters, but the effect is more pronounced for Pol IIIγ. This implies that −1PRF is not necessary for the Uup effect and that Uup can modulate the expression of both Pol IIIγ and Pol IIIτ.

Moreover, expression of all the *dnaX* constructs, with the exception of the one expressing only a short fragment of the *dnaX* sequence (*yfp*:*dnaX*_624-643aa_), led to toxicity in the bacteria, as evidenced by the reduction in growth rate compared to the condition without IPTG, where the construct is not expressed (**EV Fig. 3C**). Interestingly, the decrease in growth rate was more pronounced in the Δ*uup* strain compared to the WT and *C_uup_* strains. These observations suggest that the fusion of YFP to the *dnaX* products causes cellular toxicity, and in the Δ*uup* strain, the lack of proper expression results in even greater toxicity.

### uup-dependent regulation is independent of EF-P

Three of the Uup targets contain a proline residue within the terminal sequences that are critical for Uup-dependent expression. Furthermore, as described in the introduction, Uup has been shown, under non-physiological conditions, to overlap with the function of EF-P in *E. coli* (Chadani *et al*., 2024). A Uup homologue, YfmR, rescues mRNA translation at polyproline stretches in *B. subtilis* (Hong *et al*., 2024). The *efp* gene was initially described as essential in *E. coli* (Aoki *et al*, 1997); however, a deletion mutant of *efp* was obtained in the Keio collection (Baba *et al*, 2006) which was later used by several laboratories (Chadani *et al*., 2024; Ude *et al*, 2013).

To investigate the interplay between Uup and EF-P, we thus constructed a Δ*uup* Δ*efp* double mutant (by transduction from the Keio collection) and assessed the expression of the *yfp*:*ssb*_129-178aa_ reporter in this strain relative to the single Δ*uup* mutant. Due to the growth defect associated with *efp* deletion, all cultures were normalized to an OD_600_ of 1 prior to measuring YFP fluorescence (**Fig. 5**). As expected, YFP expression was reduced in the Δ*uup* strain compared to both wild-type and *C_uup_* strains under these conditions (**Fig. 5**). When comparing the Δ*uup* Δ*efp* strain to the Δ*efp* and Δ*efp C_uup_* strains, we observed a reduction in YFP reporter expression in the double mutant. These findings suggest that the regulatory effect of Uup on reporter expression is at least partially independent of EF-P.

**Figure 5:**
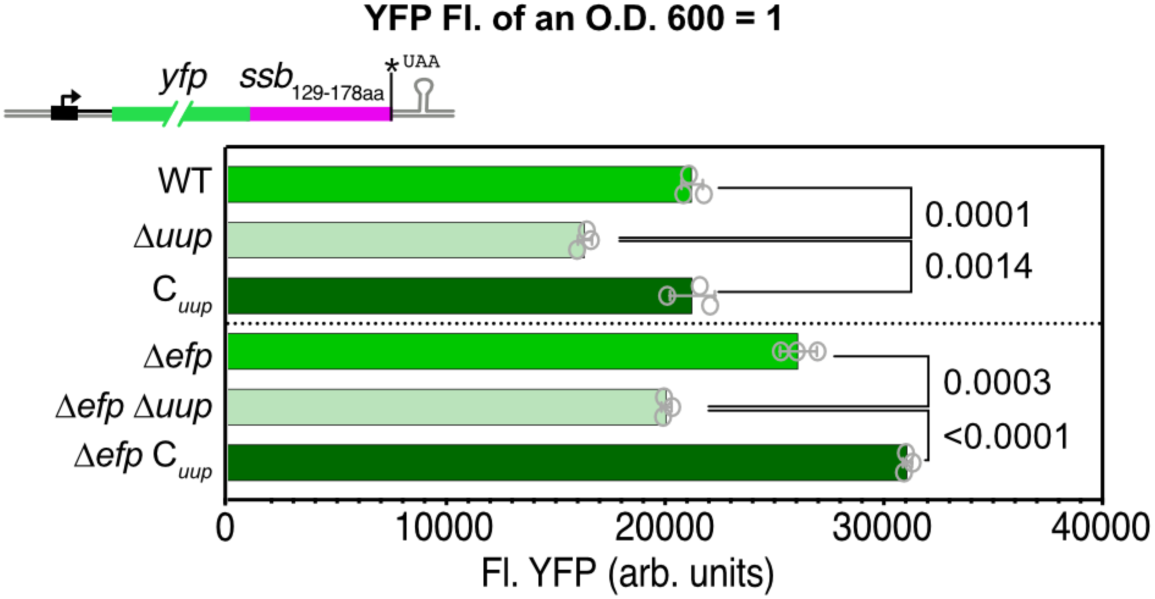
Uup-dependent expression of the *yfp*:*ssb*_129-178aa_ reporter is independent of EF-P. Overnight cultures expressing the *yfp*:*ssb*_129-178aa_ reporter in the WT, Δ*uup*, C*_uup_*, Δ*efp* or Δ*uup* Δ*efp* genetic backgrounds were normalized to an OD_600_ of 1, and YFP fluorescence was measured. Error bars represent the mean ± s.d. for triplicate experiments. *P*-values were calculated using an unpaired two-tailed *t*-test.

### Uup rescues stalled terminating ribosomes and prevents targeting to the trans-translation pathway

Ribosome stalling during or near the termination step of mRNA translation can be rescued by the trans-translation pathway (Collier *et al*, 2004; Hayes *et al*, 2002; Woolstenhulme *et al*, 2013). This pathway involves the tmRNA, encoded by the *ssrA* gene, which is mimicking a tRNA and is charged with alanine by a tRNA-synthetase. With the help of the SmpB protein and the elongation factor EF-Tu, the tmRNA-Ala binds to the A site of the stalled ribosome. EF-G then catalyzes translocation and refolding of tmRNA as an mRNA to direct the addition of a short peptide tag to the growing polypeptide. After termination of tmRNA translation, the tagged protein is recognized by proteases and degraded.

Originally, trans-translation was characterized on non-stop mRNA *i.e*. truncated mRNA lacking the 3’ part, including the stop codon, but it has been demonstrated that when ribosome stalling occurs at termination due to the presence of a terminal Pro or other specific motifs (often including Pro), trans-translation can rescue the stalled complex (Collier *et al*., 2004; Hayes *et al*., 2002; Woolstenhulme *et al*., 2013). The insertion of tmRNA into the mRNA tunnel likely involves cleavage of the mRNA present in the stalled ribosomal complex at the A site, but the cleavage mechanism is not clearly defined (Hayes & Sauer, 2003; Janssen *et al*, 2013; Sunohara *et al*, 2004). Moreover, no specific endonuclease has been identified so far for this process (Hayes & Sauer, 2003). Relatedly, downstream colliding ribosomes can induce other mRNA cleavage sites between the stalled ribosome and the collided one (Saito *et al*, 2022).

To determine whether trans-translation was involved in rescuing stalled ribosomes in the absence of Uup, we tested our constructs in two different genetic backgrounds. First, we constructed strains deleted for the *smpB* gene (Δ*smpB*), which encodes the SmpB protein necessary for the trans-translation response, in both WT and Δ*uup* backgrounds. Second, we used the *ssrAH7* strain (PHB2392), constructed by the Bouloc laboratory, where the *ssrA* gene was modified to replace part of the protease-tagging sequence with a hepta-histidine (His_7_) encoding sequence (Collier *et al*., 2004). This modified tmRNA prevents degradation of the tagged protein and makes it detectable with an anti-His antibody. We thus deleted *uup* (Δ*uup*) in this genetic background (*ssrAH7*).

The expression of the *yfp*:*ssb*_129-178aa-UGA_ reporter in the Δ*smpB* and Δ*smpB* Δ*uup* strains showed no statistically significant difference, demonstrating that the differential expression in the Δ*uup* strain depends on the trans-translation rescue system (**Fig. 6A**). Similarly, its expression in the *ssrAH7* and *ssrAH7 Δuup* strains also showed no difference. In both cases, expression of the *yfp*:*ssb*_129-178aa-UGA_ reporter resulted in reduced growth rate (toward the stationary phase) in all tested strains. However, in the *ssrAH*7 genetic background, the deletion of *uup* exacerbated it further (**EV Fig. 4A**). Western blot analysis of protein extracts from the WT, Δ*uup* and C*_uup_* strains with an anti-GFP antibody revealed a decreased signal for the reporter in the Δ*uup* strain compared to the other two strains, confirming that the fluorescent reporter correlates with protein expression. The same extracts showed no signal when probed with an anti-His antibody. For extracts from the *ssrAH7* and *ssrAH7 Δuup* strains, no difference was observable with the anti-GFP antibody, but a strong band with equivalent migration to that observed with the anti-GFP antibody appeared when probed with an anti-His antibody (**Fig. 6A**). This observation demonstrates that, in the absence of Uup, the *ssb* translation reporter is tagged by the modified tmRNA. Moreover, when we ran the *ssrAH7 Δuup* protein extract over a Ni-NTA column, a peak was detected during elution with imidazole, and fluorescence measurement for the YFP signal showed that the peak contained YFP (**Fig. 6A insert**). This demonstrates that the full YFP is synthesized. Mass spectrometry analysis of trypsin-digested peptides from the elution fraction did not allow us to identify the tmRNA-tagged sequence, likely due to impaired detection of the tagged peptide.

**Figure 6:**
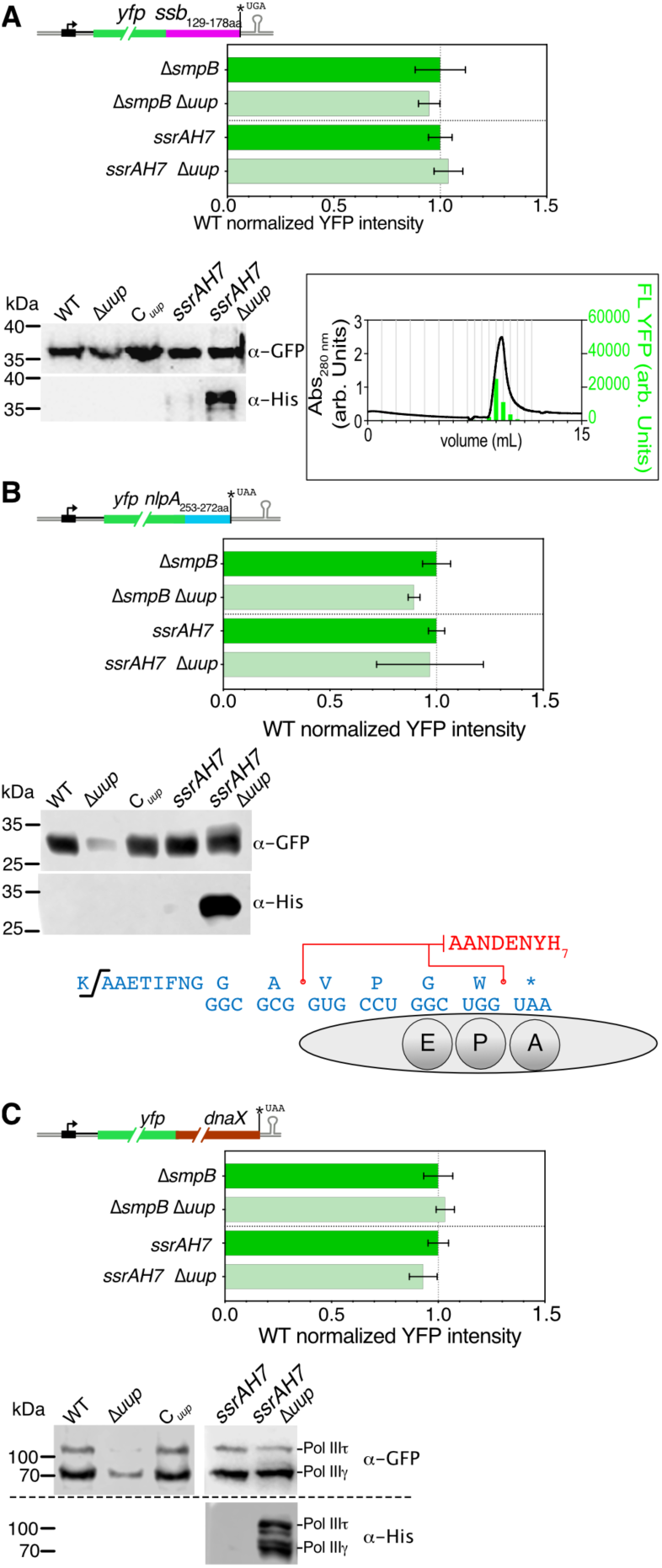
tmRNA (*ssrA*) rescues stalled ribosome in absence of Uup. Top: Histograms show fluorescence ratios normalized to the strain not deleted of *uup,* of different constructs measured in the Δ*smpB* and *ssrAH7* genetic backgrounds, with or without deletion of *uup* after 30 hours of growth in microplate (full growth curves and fluorescence are presented in **EV fig. 4**). Error bars represent mean ± s.d. from triplicate experiments, calculated as described in the methods section. Below: Western blots of protein extracts from exponential cultures of the WT, Δ*uup*, C*_uup_*, *ssrAH7* and *ssrAH7* Δ*uup* strains probed with anti-GFP or anti-His_6_ antibodies. **A)** *yfp:ssb*_129-178aa_ construct. Insert: elution profile of an Ni-NTA purification of protein extract from the *ssrAH7* Δ*uup* strain with the YFP fluorescence of the eluted fractions. **B)** *yfp:nlpA*_253-272aa_ construct. Sequence below shows the position at which the SsrAH7 tag is inserted in the protein sequence of the construct as determined by mass spectrometry. **C)** YFP reporter containing the full *dnaX* sequence encoding for both the tau (Pol III τ) and gamma subunit (Pol III γ).

For the *yfp:nlpA*_253-272aa_ construct, we conducted the same experiments. As with the *ssb* construct, no statistically significant variation in fluorescence signal was observed between the Δ*smpB* and Δ*smpB* Δ*uup* strains, nor between the *ssrAH7* and *ssrAH7 Δuup* strains (**Fig. 6B**). In all strains, expression of the *yfp:nlpA*_253-272aa_ construct induced growth stress; however, this stress was systematically increased in the Δ*uup* strain (**EV Fig. 4B**). Western blot analysis with an anti-GFP antibody showed reduced expression of the reporter only in the Δ*uup* strain compared to WT and C*_uup_* strains. In the *ssrAH7* genetic context, the anti-GFP antibody revealed similar expression between *ssrAH7 and ssrAH7 Δuup* strains but an anti-His antibody showed a strong signal only for the *ssrAH7 Δuup* strain (**Fig. 6B**), confirming that the *nlpA* construct is tagged by tmRNA in the absence of Uup. Moreover, we purified the His-tagged protein by Ni-NTA column chromatography and analyzed the trypsin-digested peptides by mass spectrometry, identifying two major peptides containing the tmRNA (*ssrAH7*) tag sequence (AANDENYH_7_). One corresponded to an insertion before the stop codon, and the second was 12 nucleotides upstream (**Fig. 6B**). The first suggests tagging when the ribosome reaches the stop codon in the A site, consistent with other results indicating that Uup helps the ribosome in terminating translation of the *nlpA* mRNA. The second may be attributed to an upstream ribosome that collided with the ribosome stalled at the translation termination step. It is known that collided ribosomes induce mRNA cleavage by the SmrB endonuclease (Saito *et al*., 2022). After cleavage, the following ribosome ends with a non-stop mRNA and is also rescued by tmRNA.

Finally, we subjected the *yfp:dnaX* construct to the same tests. With this construct, we made the same observations as with the other two. The fluorescence signal was identical between the Δ*smpB* and Δ*smpB* Δ*uup* strains, as well as between the *ssrAH7 and ssrAH7 Δuup* strains (**Fig. 6C**), confirming the involvement of trans-translation. Growth curves showed a decrease in the Δ*uup* strain in the Δ*smpB* genetic background (**EV Fig. 4C**). Western blot analysis with an anti-GFP antibody revealed reduced expression of the reporter only in the Δ*uup* strain for both protein isoforms (Pol IIIτ and Pol IIIγ) compared to the WT and C*_uup_* strains, while no difference was observed between the *ssrAH7* and *ssrAH7 Δuup* strains. The anti-His antibody revealed strong labeling for both isoforms only in the *ssrAH7 Δuup* strains. These results thus confirm that the decreased expression of *dnaX* in the absence of Uup is also related to the trans-translation system and that both protein isoforms, the one produced without −1PRF and the one produced after it, are tagged by tmRNA. This confirms the observations made with the truncated reporter (**Fig. 4C and EV Fig. 3C**), where translation termination of both isoforms appears to be altered by the absence of Uup.

### In vitro assays confirm that Uup rescues abortive termination complexes

*In vitro* translation assays (IVTA) of a *yfp:nlpA*_253-272aa_ mRNA produced *in vitro* with the T7 polymerase, were carried with or without His_6_-Uup or His_6_-Uup-EQ_2_ (a Uup mutant which binds ATP but for which the ATPase activity is inhibited) purified proteins (**Fig. 7A and EV Fig. 5**). The final amount of YFP-fusion protein was estimated by fluorescence and normalized to the control condition (no addition of Uup). Addition of His_6_-Uup increased mRNA translation by 1.6-fold, while addition of His_6_-Uup-EQ_2_ reduced it by more than 2-fold (**Fig. 7A**). We pelleted the ribosomes by ultracentrifugation at the end of each reaction and analyzed by western blot whether the YFP fusion (anti-GFP antibody) or the Uup proteins (anti-His antibody) were associated with the ribosomal pellet. In the reaction without Uup, some YFP fusion remained associated with the ribosome, and even more when His_6_-Uup-EQ_2_ was added (**Fig. 7B**). In contrast, addition of His_6_-Uup reduced the amount of YFP associated with the ribosome. These results confirm the direct action of Uup on the translating ribosome and suggest that Uup functions by improving the release of the nascent protein from the ribosome. They also show that the ATPase activity of Uup is necessary in two ways for this process: by participating in the release of the nascent protein (since the His_6_-Uup-EQ_2_ increases the amount of nascent protein associated with the ribosome) and by allowing the dissociation of Uup from the ribosome (since only His_6_-Uup-EQ_2_ is detected in the ribosomal pellet). Further work is needed to disentangle the dual ATPase activity of Uup.

**Figure 7:**
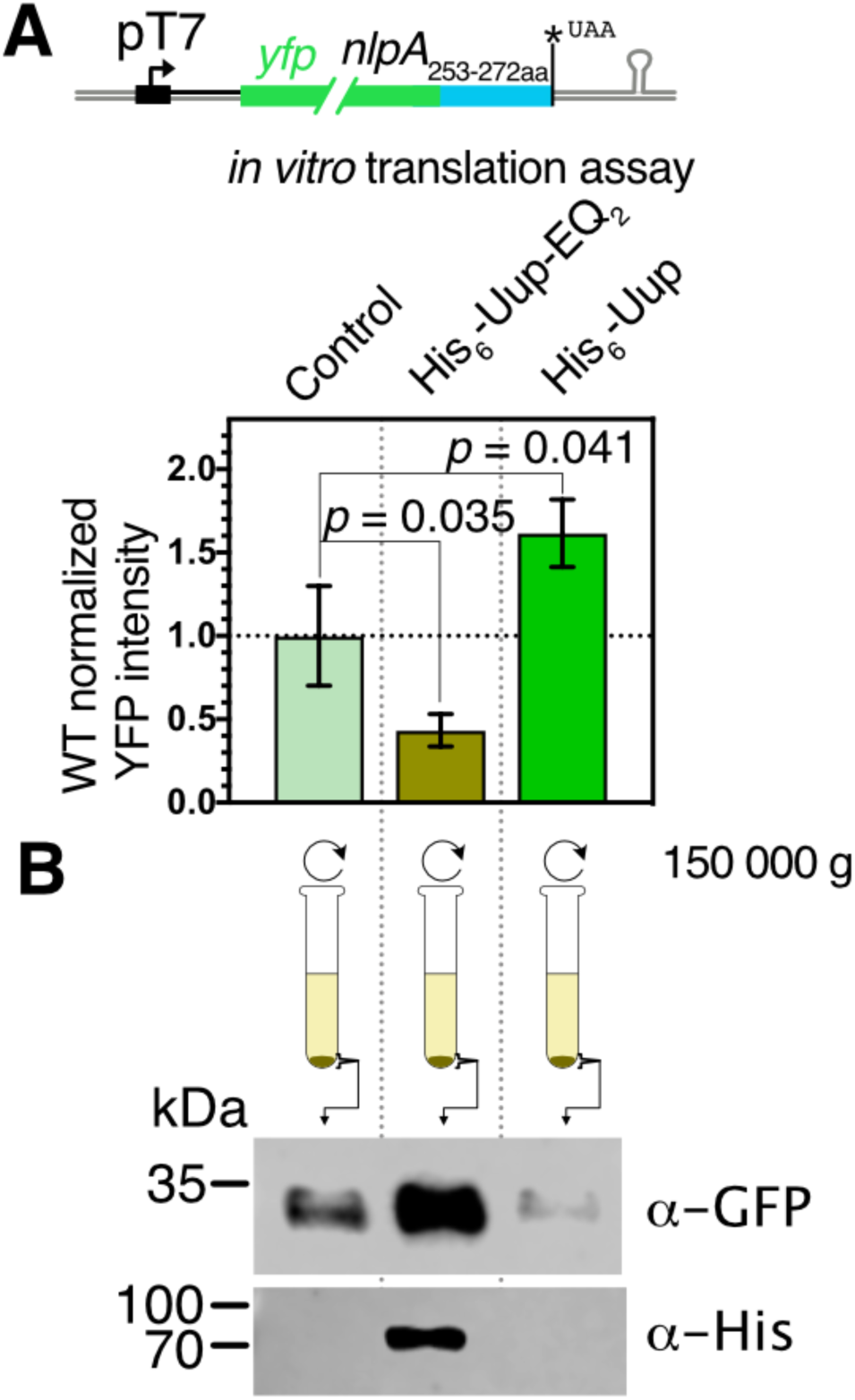
***In vitro* translation assay demonstrating the direct role of Uup in rescuing translation termination of the *yfp*:*nlpA*_253-272aa_ mRNA. A)** Minimum *in vitro* translation assay using PUREFrex 1.0 cell free protein synthesis kit (GeneFrontier Corporation) in presence of 0.5 μM of purified MRE600 ribosomes and 0.5 μM of the purified *yfp*:*nlpA*_253-272aa_ mRNA (obtained by T7 *in vitro* transcription). The assay includes the addition of the Uup purification buffer (Control), 1 μM of purified His_6_-Uup-EQ_2_ or 1 μM of purified His_6_-Uup. Reactions were incubated at 37°C and YFP fluorescence was measured after 2 h**. B)** Reactions were pelleted by ultracentrifugation to isolate ribosomes, and the pellets were analyzed by Western blot probed with anti-GFP or anti-His₆ antibodies.

### The low expression of ssb in the Δuup strain is responsible for the elongated cell phenotype

To connect our study back to our primary question about the discrepancy between the role of Uup in DNA maintenance and its role in mRNA translation, we hypothesized that the Δ*uup* phenotype related to DNA maintenance, such as the elongated cell phenotype, was due to a lack of SSB expression. To test this hypothesis, we exploited our finding that replacing the terminal Phe of SSB with Tyr alleviates its Uup dependency for expression; producing SSB protein at similar levels in both the WT and Δ*uup* strain (**Fig. 4A and EV Fig. 3A**).

Thus, we expressed the full SSB with Phe178 replaced by Tyr (variant SSB_F178Y_) in the Δ*uup* strain, on a pBAD plasmid under the control of an arabinose-inducible promoter and observed cells by microscopy at two different OD_600_ values (0.5 and 1.5) and under conditions of repression (0.4% D-glucose) or induction at different levels (0.001%, 0.01%, and 0.1% L-arabinose). Under repressive conditions, we observed the same elongated cell phenotype (**Fig. 8**) as in **Fig. 1A and EV Fig. 1A** for the Δ*uup* strain. However, with induction, the elongated cell phenotype was reduced at 0.001% arabinose, while completely absent at 0.01% and 0.1%. This result suggests that increasing SSB expression in the Δ*uup* strain can rescue the phenotype and strongly supports the idea that the phenotype induced by Uup deletion is, at least in part, due to reduced SSB expression in this strain.

**Figure 8:**
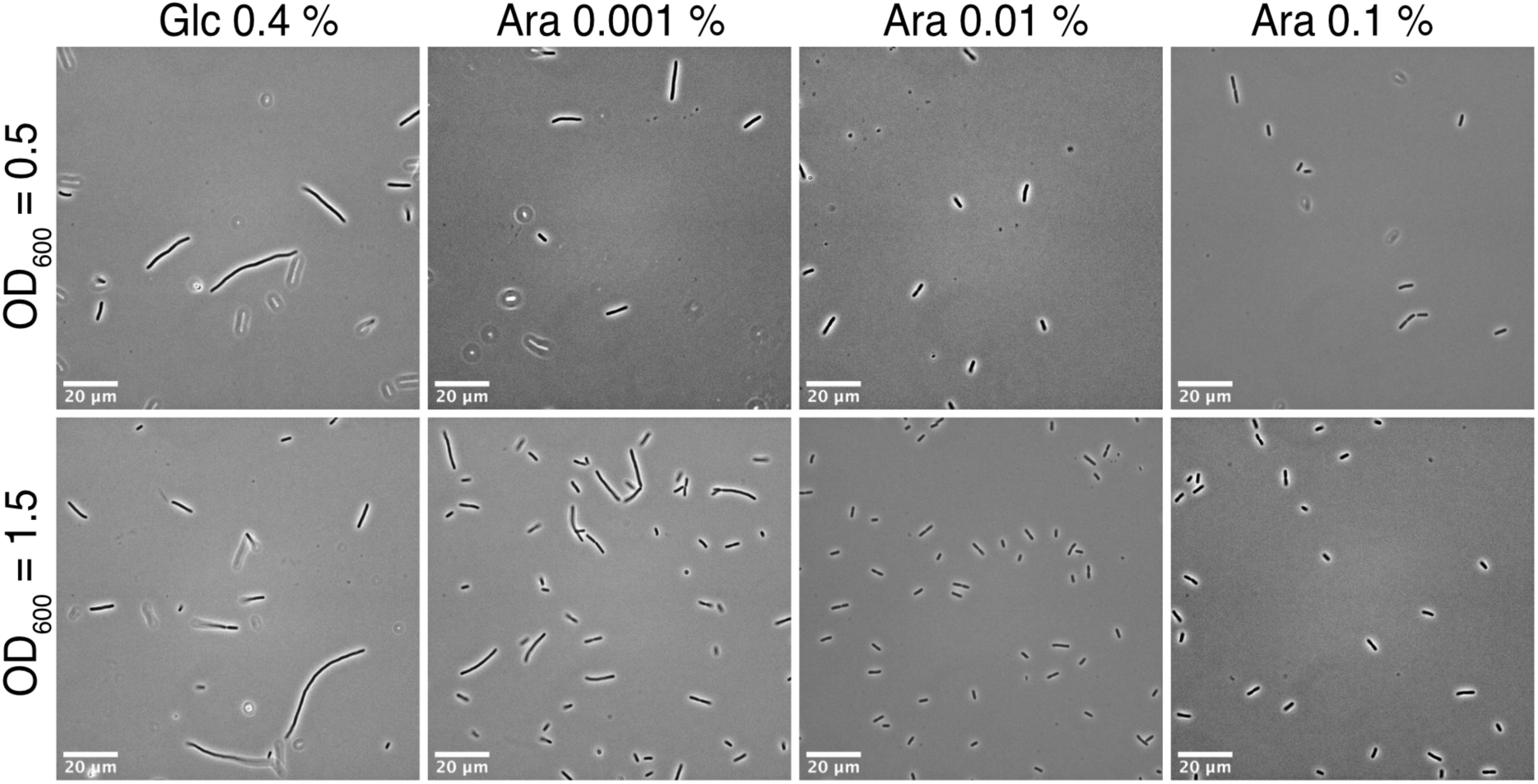
The Uup-independent SSB_F178Y_ variant rescues cell-elongation phenotype caused by Uup loss. Phase-contrast microscopy of the Δ*uup* pBAD-*ssb*_F178Y_ strain expressing SSB_F178Y_ protein (which is Uup-independent for its expression) upon L-arabinose induction. Cells were harvested at an OD₆₀₀ of 0.5 or 1.5 after culture in LB medium supplemented with 0.4% D-glucose (Glc) for repression or 0.001%, 0.01%, and 0.1% L-arabinose (Ara) for varying levels of induction.

## Discussion

From the observation that polyproline stretches impair mRNA translation primarily due to ribosome stalling, and the concurrent discovery of the dedicated factor EF-P that resolves this issue, it became evident that other mRNA translation contexts could also pose challenges for the ribosome. Indeed, similarly to the Context-Dependent Ribosomal Stalling (CDRS) that are associated with stalling induced by antibiotics targeting translation elongation (Vázquez-Laslop & Mankin, 2018), CDRS can also occur independently of small molecules binding to the ribosome. It was shown that antibiotic-independent CDRS mechanisms, such as repeats of acidic residues within the first 30 amino acids of an elongating protein, arise not only from ribosome stalling but also from ribosome dissociation. In such context, the dedicated translation factor EttA prevents dissociation or stalling (Chadani *et al*., 2024; Ousalem *et al*., 2024). Another form of CDRS occurs during translation termination, where combination of specific residues at the C-terminus can stall the termination complex by inhibiting the hydrolysis of the nascent peptide ester bond from the tRNA, a reaction normally catalyzed by release factors RF-1 or RF-2 (Collier *et al*., 2004; Hayes *et al*., 2002; Woolstenhulme *et al*., 2013). The C-terminal sequence involved in such a stalling are varied but often contain proline or tryptophan residues (Collier *et al*., 2004; Hayes *et al*., 2002; Woolstenhulme *et al*., 2013). Such stalling events often trigger the tmRNA rescue response, wherein tmRNA binds to the A site of the stalled ribosome at the stop codon. Following transfer of the nascent peptide to the alanine carried by tmRNA, the ribosome translates the tmRNA coding sequence, tagging the nascent peptide for recognition and degradation by the degradosome.

To date, no translation factor capable of rescuing this type of ribosome stalling had been identified. Herein, we present an extensive characterization of the function of Uup in rescuing stalled translation complexes at termination. After identifying several phenotypes associated with DNA maintenance in the Δ*uup* strain (**Fig. 1**), we employed an unbiased global approach to identify proteins whose expression was reduced in the Δ*uup* strain (**Fig. 3**). Among the identified proteins, SSB and the Pol IIIτ and Pol IIIγ subunits encoded by the *dnaX* gene, are involved in DNA maintenance, while another, NlpA, is a lipoprotein. Using fluorescence translation reporters, we demonstrated that, for all four proteins, the effect of Uup on their expression resulted from alleviating an mRNA translation bottleneck at the termination step (**Fig. 4**), and that this effect was dependent on the nature of the stop codon. We further demonstrated that this regulation is independent of EF-P (**Fig. 5**) and confirmed ribosome stalling through two distinct approaches. First, we showed that the absence of Uup targets the stalled complex to the tmRNA rescue pathway (**Fig. 6**), as expected for stalled complexes at or near termination. For the target NlpA, we identified that tmRNA rescue resulted in the insertion of the tmRNA tag either immediately before the stop codon or four codons upstream, consistent with a ribosome stalled with a stop codon in the A site or due to a collision with a trailing ribosome (**Fig. 6B**). Second, using an IVTA, we observed that the addition of Uup increased the expression of one target, NlpA (**Fig. 7A**). In the absence of Uup, the nascent peptide remained associated with ribosomes. Notably, the addition of the ATPase-deficient mutant Uup-EQ_2_ further reduced NlpA expression and increased the amount of nascent peptide associated with the ribosome while the Uup-EQ_2_ protein remained bound to the ribosome (**Fig. 7B**). These observations strongly suggest that the ATPase activity of Uup is required to resolve stalling triggered by the lack of hydrolysis of the tRNA-nascent peptide.

Moreover, our study resolves a long-standing conundrum regarding Uup function. It was originally proposed that Uup is directly involved in transposon release through direct interaction with DNA, as well as in modulating ribosome activity (see introduction). Here, we demonstrate that the involvement of Uup in DNA maintenance is a secondary effect, most likely due to the reduced expression of SSB and the Pol IIIτ and Pol IIIγ subunits. We further show that ectopic expression of a SSB variant which is no longer dependent on Uup for its expression (SSB_F178Y_), can rescue the elongated cell phenotype induced by genomic instability in the Δ*uup* strain (**Fig. 8**). Interestingly, *ssb* gene was also targeted in the precise transposon release screen that originally identified *uup*, as the other mutations identified in that study were in the *ssb* gene (Reddy & Gowrishankar, 1997). Therefore, the primary function of Uup is to resolve stalled translation termination complexes, and this action regulates the expression of key proteins involved in DNA maintenance. It is possible that Uup connects DNA replication with translation, but further studies will be needed to test this hypothesis.

Finally, we propose a model (**Fig. 9**) illustrating the dual mechanisms by which bacterial cells resolve stalled ribosome complexes during translation termination. Stress-generating aa sequences induce ribosomal stalling through a bifactor effect: *i*) improper structural conformation of aa chain within the ribosomal exit tunnel and *ii*) specific recently incorporated residues that prevent hydrolysis of the aa-tRNA ester bond by the RF due to the improper conformation of aa chain within the ribosomal exit tunnel. The left panel depicts the Uup rescue pathway, in which the Uup protein, in an ATP-dependent manner, binds to the stalled ribosome. Its ATPase activity, through an uncharacterized mechanism, facilitates hydrolysis of the ester bond possibly in coordination with the RF, after which Uup dissociates from the complex. This process ensures the optimal synthesis of essential proteins such as NlpA, SSB, Pol IIIγ, and Pol IIIτ. In the absence of Uup (Δ*uup*, right panel), the tmRNA rescue pathway is activated. Here, the *trans*-translation system, comprising SmpB and tmRNA (*ssrA*), tags the stalled nascent polypeptide, marking it for degradation. This pathway ensures the clearance of non-functional or potentially harmful proteins, thereby maintaining cellular homeostasis. The model highlights the critical role of Uup in resolving stalled translation complexes and its indirect impact on DNA maintenance through the regulation of key protein synthesis.

**Figure 9:**
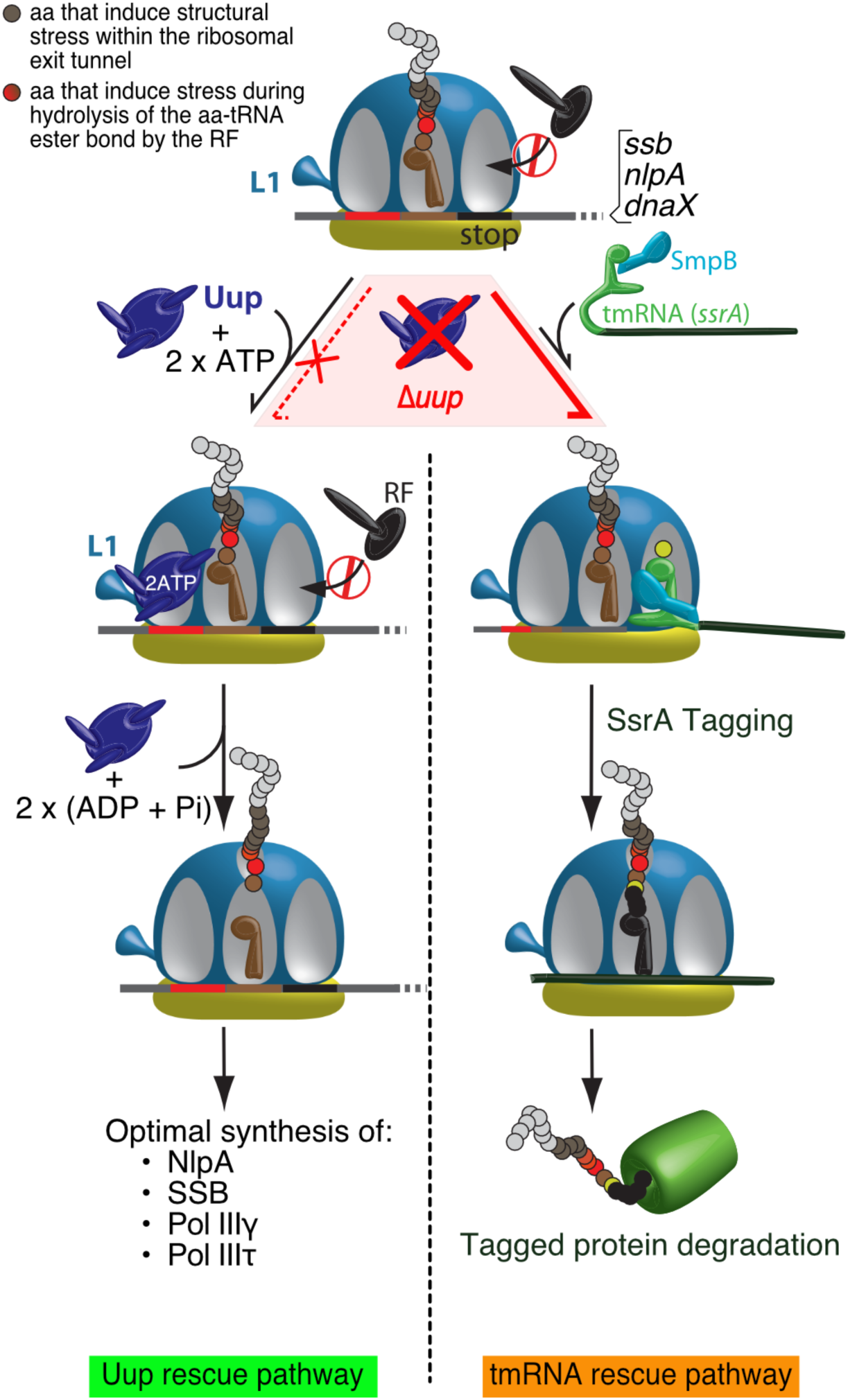
Model of Uup- and tmRNA-mediated rescue pathways for stalled ribosome complexes. Schematic representation of the two distinct rescue pathways for ribosome stalling during translation termination. In the presence of Uup (left panel), ATP hydrolysis by Uup resolves stalled ribosome complexes, enabling optimal synthesis of the proteins NlpA, SSB, Pol IIIγ, and Pol IIIτ. In the absence of Uup (Δ*uup*, right panel), the *trans*-translation system, involving SmpB and tmRNA (*ssrA*), tags the stalled nascent polypeptide, leading to its degradation. Ribosomal stalling is induced by a bifactor effect: *i*) the last amino acid within the ribosomal exit tunnel (brown circles) induces improper structural positioning of the peptide, *ii*) specific terminal residues (red circles), due to improper conformation, prevent hydrolysis of the aa-tRNA ester bond by the release factor (RF).

## Methods

### Culture conditions

The bacterial strains used in this study are listed in **Supplementary Table 1**. Bacteria were routinely cultured in Luria-Bertani Miller broth (LB-Difco) with or without addition of appropriate antibiotics (ampicillin at 100 µg·ml^−1^, kanamycin at 25 µg·ml^−1^, chloramphenicol at 20 µg·ml^−1^) at 37 °C with 180 rpm agitation. Preinocula of the cultures were set up in 3 ml LB and cultured overnight and were used in 1:1000 dilution. Growth of the cultures was determined by measurement of OD_600_ using UviLine 9100 (Secomam). 1mM Isopropyl β-D-1-thiogalactopyranoside (IPTG) was added to the media for induction of pMMB-based constructs. For induction of pBAD-based constructs 0.1% (w/v) L-arabinose was added to the media. Expression of pBAD plasmid was repressed by addition of 0.4% (w/v) of ß-D-glucose in preinocula cultures. For measurement of the growth and expression of the pMMB-based constructs and Uup:YFP induction, a CLARIOstar plate reader (BMG Labtech) was used. The cultures were set up in 96-well Falcon microplates in 200 μl of LB with appropriate antibiotics and inducers, inoculated with 2μl of preinocula and topped with 60 μl (or 40 μl for YFP-SSB constructs) of mineral oil (Merck) to prevent evaporation. Every 30 min measurements of OD_600_ and YFP emission were collected under 600 rpm orbital shaking. YFP was excited at 497-15 nm, and the fluorescence was collected in the range of 540-20 nm.

### Strains construction

All Δ*smpB*, Δ*efp*, backgrounds as well as the Δ*uup ssrAH7* strains were produced by P1 transduction. The parental strains and the donor strains for the transduction are specified in **Supplementary Table 1**. The strains were validated by sequencing with primers listed in Supplementary Table 3. The *uup::yfp* strain was constructed by inserting a *yfp* fragment generated by PCR described in **Supplementary Table 3**. The PCR product was introduced into MG1655 cells harboring the pKD46 plasmid (which expresses the λ-Red recombinase and Gam proteins) via electroporation, followed by selection for chloramphenicol resistant clones (Datsenko & Wanner, 2000).

### Construction of plasmids

Plasmids and primers used in this study are listed in **Supplementary Tables 2 and 3**. All the newly constructed plasmids were cloned in DH5α strain and then moved to MG1655 backgrounds by TSS transformation (Chung *et al*, 1989). The plasmids were extracted using NucleoSpin Plasmid EasyPure kit (MACHEREY-NAGEL) according to the manufacturer’s instructions.

The pMMBpLacO_TBT_*yfp* plasmid used as a backbone for target validation was derived from the IPTG inducible low-copy plasmid pMMBpLlacO-1-67EH (Fostier *et al*., 2023). The TBT YFP variant was chosen as a reporter protein due to its increased detectability (Lipońska *et al*, 2025). The constructs were produced by amplification of the insert and vector fragments by PCR with CloneAmp HiFi PCR Premix (Takara Bio). Primers used for the amplification of the fragments contained overlap sequences, which allowed for assembly of the constructs with NebBuilder HiFi DNA Assembly Master Mix (New England Biolabs). The amino acid, stop codon and synonymous codon substitutions were introduced on the overhang regions of the primers that harboured sequences homologous to the plasmid sequence. This allowed for assembly of the insert and vector fragments or recircularisation of the plasmid with the introduced mutation using the same strategy. Production of pBADØ fragment used in construction of pBAD-*ssb_F_178_Y_*UAA* was an exception: instead of PCR amplification, a digestion with NcoI (New England Biolabs) was performed to linearise the plasmid. pMMB-*yfp:nlpA253-272aa_PGW_270-272__synonymous_codon_*UAA* construct introduced changes in pMMB-*yfp:nlpA253-272aa_*UAA* P_270_ from CCT to CCG and G_271_ from GGC to GGG. This resulted in production of the same residues but with a different coding sequence. All of the constructions were validated by sequencing with NightXpress Mix2Seq Kit (Eurofins).

### Survival drop assay and Uup:YFP induction

For survival drop assay duplicate of MG1655 (WT), Δ*uup* and C_uup_ strains (Supplementary Table 1) cultures were set up in LB medium from saturated preinocula. For Uup induction experiment triplicate of *uup:yfp* and MG1655 (WT) strains were cultured in LB medium from overnight preinocula. In both cases the cultures were grown at 37 °C with 180 rpm shaking OD_600_ = 0.5. Five ml of each culture replicate was transferred into a petri-dish and was exposed to 450 000 μJ / cm^2^ of Ultraviolet (UV) 254 nm in UVP HL-2000 Hybrilinker (Fischer Scientific). For survival drop assay experiment, the UV-exposed cultures and control non-exposed cultures were serially diluted and 5 μl drops were plated on pre-dried LB plates. The plates were incubated overnight at 37 °C. For the Uup induction experiment, 200 μl of the exposed and non-exposed cultures topped with 60 μl of mineral oil were incubated in CLARIOstar plate reader (BMG Labtech) and the YFP fluorescence was tracked as described above.

### Microscopy

Most of the samples were collected at OD_600_ = 0.7 excepted for the experience with the SSB_F178Y_ variant (**Fig. 8**) where samples were collected at OD_600_ = 0.5 or 1.5. For MG1655 (WT) and *Δuup* samples, 0.1 μg of 4′, 6-Diamidino-2-phenylindole dihydrochloride (DAPI) (Merck) was added to 1ml of each culture and the aliquots were incubated at 37 °C with shaking for 20 min. For all the strains, 1 μl of a sample was placed on a glass microscopy slide (Knittel Glass) and covered with a cover slip (Menzel-Gläser). The samples were imaged using a Zeiss Axio Observer Z1 microscope with an ORCA-Flash40 (Hamamatsu) camera. The microscopy was performed under oil immersion with Plan-Apochromat 100x objective (Zeiss) and the images were acquired using ZEN 3.8 software in Phase (WT, Δ*uup*, Δ*uup* pBADØ and Δ*uup* pBAD-*ssbF_178_Y_*UAA*), Brightfield (WT p*uup*-EQ_2_) and DAPI fluorescence (WT and Δ*uup*). Cell length was determined with ImageJ, MicrobeJ 5.13l using a default set up for length calculation (Ducret *et al*, 2016).

### Protein and peptide purification

His_6_-Uup and His_6_-UupEQ_2_ were purified as previously described (Ousalem *et al*., 2023) after expression in the LOBSTR strain (Andersen *et al*, 2013). For purification of *ssrAH7*-targetted peptides of the Uup targets, pMMB-*yfp:nlpA253-272aa_*UAA* and pMMB-*yfp:ssb129-178aa_*UGA* (**Supplementary Table 2**) were expressed in *Δuup ssrAH7* (**Supplementary Table 1**) background until OD_600_ = 1 at 37 °C with 180 rpm. The cultures were centrifuged at 5000 g for 10 min at 4 °C. Pellets were resuspended in 30 ml of 5 μM imidazole protein purification buffer (20 mM Tris base pH 7.5, 300 mM NaCl, 10 % glycerol, 5 mM imidazole pH 7.5, 2 mM β-mercaptoethanol) with addition of 1 mg/ml lysozyme, 50 μl protease inhibitor cocktail (Sigma–Aldrich) and incubated on ice for 30 min. The lysis suspension was passed 6 times through a cell disrupter (Constant Systems Ltd) at 2.5 kbar pressure. The lysate was centrifuged at 24000 g for 20 min at 4 °C and the supernatant was loaded onto 1ml Protino Ni-NTA column (MACHEREY-NAGEL). The peptides were eluted with use of ÄKTA Start (Cytiva) with stepwise increases of the imidazole concentration in the protein purification buffer (20 mM, 40 mM and 500 mM). Collected fractions were migrated on 12.5% acrylamide SDS-PAGE gel and immunoblotted for the presence of YFP (anti-GFP) and SsrAH7 tag (anti-His_6_) as described below.

### Polyribosome fractionation

For the ribosomal gradient analysis of MG1655 expressing the ABC-F EQ_2_, saturated precultures of *E. coli* MG1655 transformed with pBAD-Ø, pBAD-*His_6_-ettA*-EQ_2,_ pBAD-*His_6_-uup*-EQ_2_ (**Supplementary Table 2**) were grown overnight at 37 °C under vigorous shaking in LB, 100 µg.ml^−1^ ampicillin, 0.4 % (w/v) β-D-Glucose. A medium of 100 ml LB medium, 100 µg.ml^−1^ ampicillin, 0.1% (w/v) L-arabinose was inoculated with the precultures, and cells were grown at 37 °C and 180 rpm. The OD_600_ measurements were taken every 30 min and the cultures were collected when the growth inhibition caused by expression of Uup-EQ_2_ was observed. For both experiments the cultures were centrifuged at 6000 rpm for 5 min at 4 °C Supernatant was discarded and cells were resuspended in 500 µl of lysis buffer (20 mM Tris pH 8, 20 mM MgOac, 100 mM NH_4_(OAc), 2 mM β-mercaptoethanol, 1 mg.ml^−1^ lysozyme, 1μl DNAse RNAse-free (ZYMO) and incubated for 2 min on ice. Then, 15 µl of sodium deoxycholate 10 % (w/v) and 15 µl BriiJ 58 10 % (w/v) were added and the samples were centrifuged for 30 min at 15 300 rpm at 4 °C. Supernatants were isolated and the total RNA concentration was quantified. The total RNA of each sample loaded onto 10-40 % (w/v) sucrose gradients made in Buffer A (20 mM Tris pH 8, 20 mM MgOAc, 100 mM NH_4_(OAc), 2 mM β-mercaptoethanol) was standardized to 1 mg. The sucrose gradients were centrifuged 2 h at 40 000 rpm at 4 °C for 2 h in the SW41 rotor (Beckman Coulter). Samples were fractionated and collected using Biocomp Piston Gradient Fractionator (BioComp Instruments), absorbance being measured at 254 nm. The fractions were collected and subjected to immunoblotting analysis.

### Immunoblotting

The samples used in immunoblotting experiments were standardized to OD_600_ = 1 by pelleting appropriate sample volume and resuspending the pellets in 100 μl 1x Laemmli sample buffer (Bio-Rad). Samples were heated at 70 °C for 15 min and migrated on 12.5% acrylamide SDS-PAGE gels at 90V alongside PageRuler Prestained Protein Ladder (Thermo Fisher Scientific). Separated proteins were transferred onto an ethanol-activated Immun-Blot PVDF membranes (Bio-Rad) at 90 V for 90 min. The membranes were blocked in PBS (Phosphate buffered saline) with 5 % (w/v) skimmed milk for 1 h at room temperature or overnight at 4 °C. The membranes were thoroughly washed with 1 % PBS and incubated with primary mouse anti-His_6_ (Covalab) or rabbit anti-GFP (Invitrogen) antibody at 1:3000 dilution in 1% PBS 0.1% (v/v) Tween-20 for 2 h. After washing in PBS-0.1% tween, membranes were incubated with a secondary HRP-conjugated anti-mouse IgG (for primary anti-His_6_) or HRP-conjugated anti-rabbit IgG (for primary anti-GFP) antibody (Covalab) at 1:20000 dilution in 1 % PBS 0.1% Tween-20. The membranes were revealed using Clarity Western ECL Substrate (Bio-Rad) on Odyssay Fc Imaging System (LI-COR).

### RNA-seq and proteomic experiment

Aliquotes of 10 ml of LB media were inoculated with MG1655 (WT) and Δ*uup* (**Supplementary Table 1**) preinocula cultivated overnight at 37 °C with 180 rpm shaking. The cultures were set up in triplicate for each of the strains. The cultures were grown at 37 °C until OD_600_ = 2 with 180 rpm shaking.

For the RNA-seq 1ml from each replicate was pulled together (3ml total) for each strain. Six ml of DNAprotect (Qiagen) was added and the samples were centrifuged at 5000 g for 5 min at 4 °C. Supernatant was discarded and the pellets were stored at −80 °C until RNA extraction. The RNA was extracted with the Direct-zol RNA MiniPrep Kit (ZYMO RESEARCH) and eluted in Molecular grade RNase and DNase free water (US Biological Life Sciences). The quality of the RNA was checked using Qiaxcel (Qiagen) with RNA high sensitivity kit (Qiagen). All samples were standardised to 1μg in 20 μl total volume. The construction of the cDNA library and the deep sequencing were performed by the iGenSeq platform at Institut du Cerveau. RNA library preparation was realised following manufacturer’s recommendations (Illumina Stranded Total RNA Prep, Ligation with Ribo-Zero Plus). Final samples pooled library preps were sequenced on Nextseq2000 ILLUMINA with P1-300 cartridge (2*100Millions of 150 bases reads), corresponding to 2×8Millions of reads per sample after demultiplexing. Sequences were aligned with Bowtie2 (paired) and counted with FeatureCounts with use of the Galaxy server. FPKM (fragments per kilobase of transcript per million fragments mapped) values were determined for both strains.

For proteomic analysis the individual samples were adjusted to equivalent of OD_600_ = 1 of cell culture and centrifuged at 5000 g for 10 min at 4 °C. Pellets were resuspended in 200 μl of lysis buffer (250 mM Tris base pH 6.8, 2 % (v/v) SDS, 10 % (v/v) glycerol) and heat-treated at 70 °C for 15 min. The concentration of the proteins was determined by Pierce BCA Protein Assay (ThemoFisherScientific). Concentration of the samples was standardised to 30 μg in 15 μl of Molecular grade RNase and DNase free water (US Biological Life Sciences). Blue Coomassie R-250 at 0.05% (w/v) and β-mercaptoethanol at 5 % (v/v) were added and the samples were migrated on 12.5 % acrylamide SDS-PAGE for approximately 1cm. Gels were stained with Blue Coomassie R-250 solution. The part of lane containing proteins were excised and subjected to a manual in-gel digestion with modified porcine trypsin (Trypsin Gold, Promega) as previously described in (Ousalem *et al*., 2024). Briefly, after destaining, bands were subjected to a 30 minutes reduction step at 56°C using 10 mM dithiothreitol in 50 mM ammonium bicarbonate (AMBIC) prior to a 1-hour cysteine alkylation step using 50 mM iodoacetamide in 50 mM AMBIC. Both reactions were performed in the dark. After dehydration under vacuum, bands were re-swollen with 500 ng of trypsin in 200 µl 50 mM AMBIC and proteins were digested overnight at 37 °C. Peptides present in gel pieces were extracted with 1% (v/v) trifluoroacetic acid and dried in a vacuum concentrator. Then, peptide mixtures were solubilized in 150 µl of solvent A (0.1% (v/v) formic acid in 3% (v/v) acetonitrile) to a final concentration of 200 ng/µl. Mass spectrometry analyses were performed on a Q-Exactive Plus hybrid quadripole-orbitrap mass spectrometer (Thermo Fisher, San José, CA, USA) coupled to an Easy 1000 reverse phase nano-flow LC system (Proxeon) using the Easy nano-electrospray ion source (Thermo Fisher). Peptide mixtures were analyzed in triplicate. Five µl of peptide mixtures were loaded onto an Acclaim PepMap™ precolumn (75 µm x 2 cm, 3 µm, 100 Å; Thermo Scientific) equilibrated in solvent A and separated at a constant flow rate of 250 nl/min on a PepMap™ RSLC C18 Easy-Spray column (75 µm x 50 cm, 2 µm, 100 Å; Thermo Scientific) with a 90 min gradient (0 to 20% B solvent (0.1% (v/v) formic acid in acetonitrile) in 70 min and 20 to 37% B solvent in 20 min). Data acquisition was performed in positive and data-dependent Top10 modes. Full scan MS spectra (mass range m/z 400-1800) were acquired in profile mode with a resolution of 70,000 (at m/z 200) and MS/MS spectra were acquired in centroid mode at a resolution of 17,500 (at m/z 200). Raw data were processed as previously described in Ousalem *et al*., 2024 using the MaxQuant software package (http://www.maxquant.org, version 1.6.0.13) (Tyanova *et al*, 2016).

For the ssrAH7 tag identification (**Fig. 6**), the protein bands at the expected weight and immunodetected by both the anti-GFP and anti-His antibodies from the purified ssrAH7-tagged YFP:NlpA253-272aa_*UAA product were processed as described above, except that the peptide mixtures were solubilized in 20 µl of solvent A. Mass spectrometry analysis was performed as described above, using the Uniprot *E. coli* K12 (release: 05/11/2024) library with the addition of 10 predicted ssrAH7-modified YFP:NlpA253-271 peptides, in combination with a homemade contaminants database.

### in vitro transcription and translation

The DNA template for generation of mRNA used in *in vitro* translation assays was produced from a purified pMMB-*yfp:nlpA253-272aa_*UAA* plasmid (**Supplementary Table 2**) with CloneAmp HiFi PCR Premix (Takara Bio). The primers used (**Supplementary Table 3**) have a homology for the backbone of the construct and the forward primer introduces the T7 polymerase promoter sequence (5’-GCGAATTAATACGACTCACTATAGGG-3’) upstream of the *yfp* sequence. *in vitro* mRNA synthesis was performed using T7 RiboMAX Large Scale RNA Production System kit (Promega). DNA template was removed from the reaction by digestion with DNAse RQ1. The mRNA was purified using TRIzol (Thermo Fisher Scientific) and Direct-zol RNA Miniprep kit (ZYMO RESEARCH) according to manufacturer’s instructions. mRNA was eluted with RNA Storage Solution (Ambion), aliquoted and stored at −80 °C.

PUREFrex 1.0 cell free protein synthesis kit (GeneFrontier Corporation) was used for *in vitro* translation assay. Solution I (containing amino acids, NTPs, tRNAs and substrates for the translation enzymes) and Solution II (translation factors in 30% glycerol buffer) from the PUREFrex 1.0 kit were used according to manufacturer’s instructions. Solution III containing ribosomes was substituted with 0.5 μM of MRE600 ribosomes purified in the laboratory (Ousalem *et al*., 2024). The purified mRNA at 0.5 μM was added to the reaction with or without supplementation with 1 μM of purified His_6_-Uup or His_6_-Uup EQ_2_. The volume of the reaction was brought up to 10 μl with Molecular grade RNase and DNase free water (US Biological Life Sciences). The reaction was incubated for 2 h at 37 °C without shaking in CLARIOstar plate reader (BMG Labtech). Translation of the mRNA was monitored every 2 min by tracking the YFP fluorescence. All *in vitro* translation reactions were set up in a triplicate. To determine the association of peptides produced from the pMMB-*yfp:nlpA253-272aa_*UAA* with ribosomes, 8 μl of each *in vitro* translation replicate was collected after 2 h of incubation. The replicates were combined and resuspended in 126 μl Buffer A (20 mM Tris pH 8, 20 mM MgOAc, 100 mM NH_4_(OAc)) and the samples were centrifuged for 20 min at 100 000 rpm at 4 °C. Pellets were resuspended in 100 μl 1x Laemmli (Bio-Rad) and subjected to immunoblotting with mouse anti-His_6_ (Covalab) and rabbit anti-GFP (Invitrogen) antibodies as described above.

### Statistics and reproducibility

All the experiments presented have been reproduced at least twice with the same results. Statistical details can be found in the figure legends. Data were plotted using Prism 8 (GraphPad). Determination of the fluorescence ratio were done as previously described in Ousalem et al., 2024 (Ousalem *et al*., 2024) with an exception that the average for the WT, Δ*uup* and C*_uup_* strains was corrected by subtracting the average of the FluoN of the WT IPTG-uninduced strain to calculate the corrected normalized fluorescence (FluoNC).

**Table 1.**
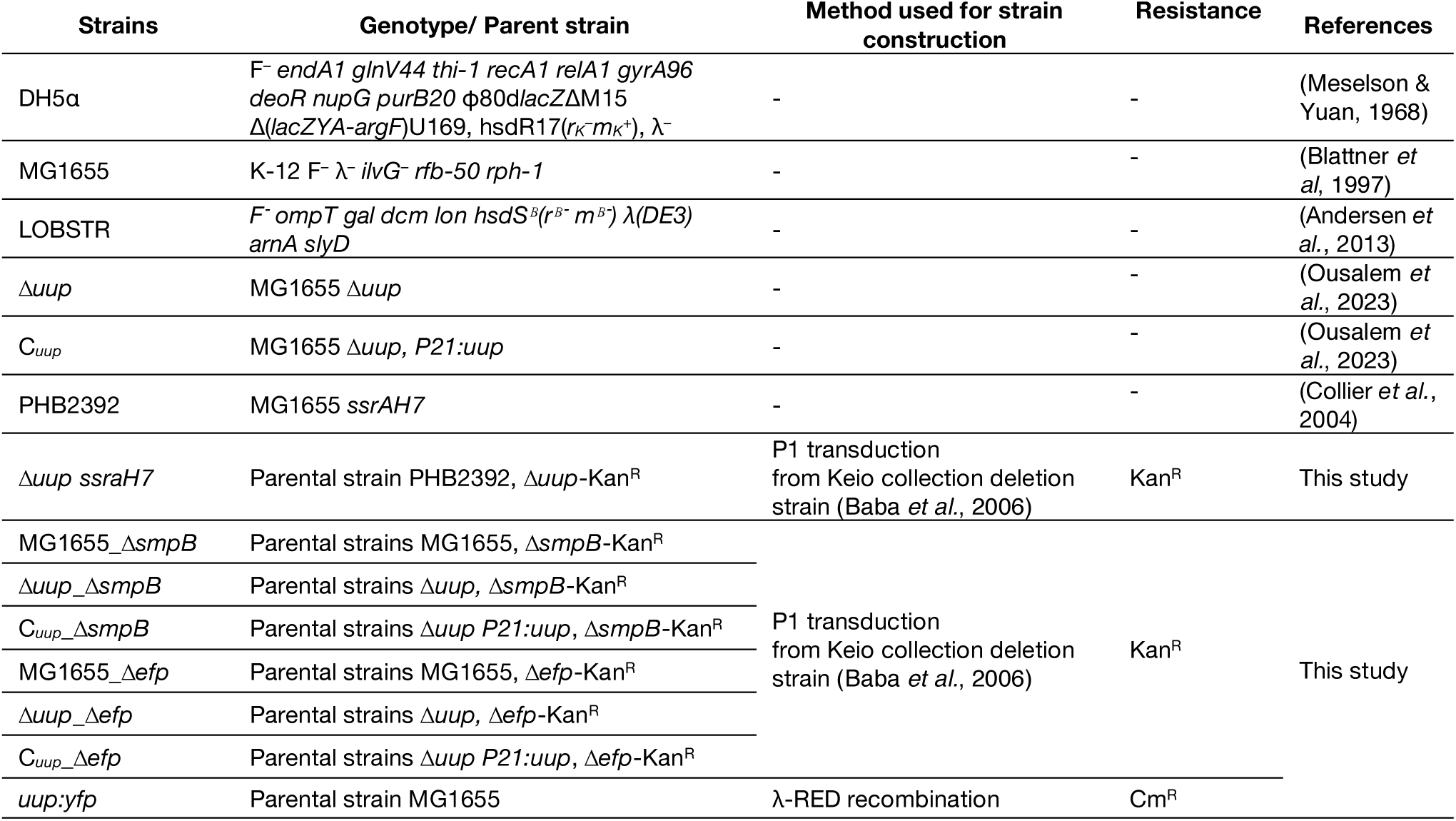
Bacterial strains.

**Table 2.** Plasmids.

| Plasmids | Selection | References |
| --- | --- | --- |
| pMMBpLacO_TBT_--yfp | Amp | This study |
| pBAD-Ø (Myc 6His) | Amp | Invitrogen |
| pBAD-*His-ettA_EQ* | Amp | (Boel *et al.*, 2014) |
| pBAD-*His-uup_EQ* | Amp | (Ousalem *et al.*, 2023) |
| pBAD-*His-uup* | Amp | (Ousalem *et al.*, 2023) |
| pKD46 | Amp | (Datsenko & Wanner, 2000) |
| pMMB-*ssb129-178aa:yfp* | Amp | This study |
| pMMB-*yfp:ssb129-178aa_\*UAA* | Amp | This study |
| pMMB-*yfp:ssb161-178aa_\*UAA* | Amp | This study |
| pMMB-*yfp:ssb129-176aa_\*UAA* | Amp | This study |
| pMMB-*yfp:ssb129-178aa_P_G_\*UAA* | Amp | This study |
| pMMB-*yfp:ssb129-178aa_F_Y_\*UAA* | Amp | This study |
| pMMB-*yfp:ssb129-178aa_PF_GG_\*UAA* | Amp | This study |
| pMMB-*yfp:ssb129-178aa_\*UGA* | Amp | This study |
| pMMB-*yfp:ssb129-178aa_\*UAG* | Amp | This study |
| pMMB-*yfp:nlpA_\*UAA* | Amp | This study |
| pMMB-*yfp:nlpA253-272aa_\*UAA* | Amp | This study |
| pMMB-*yfp:nlpA253-269aa_\*UAA* | Amp | This study |
| pMMB-*yfp:nlpA253-272aa_P_G_\*UAA* | Amp | This study |
| pMMB-*yfp:nlpA253-272aa_W_G_\*UAA* | Amp | This study |
| pMMB-*yfp:nlpA253-272aa_W_Y_\*UAA* | Amp | This study |
| pMMB-*yfp:nlpA253-272aa_PGW_synonymous_codon_\*UAA* | Amp | This study |
| pMMB-*yfp:nlpA253-272aa_\*UGA* | Amp | This study |
| pMMB-*yfp:nlpA253-272aa_\*UAG* | Amp | This study |
| pMMB-*yfp:dnaX_\*UAA* | Amp | This study |
| pMMB-*yfp:dnaX_\*UGA* | Amp | This study |
| pMMB-*yfp:dnaX_\*UAG* | Amp | This study |
| pMMB-*yfp:dnaX624-643_\*UAA* | Amp | This study |
| pMMB-*yfp:dnaX624-643_\*UGA* | Amp | This study |
| pMMB-*yfp:dnaXy_isoform_\*UAG* | Amp | This study |
| pMMB-*yfp:dnaXy_isoform_\*UGA* | Amp | This study |
| pBAD-*ssb_F_Y_\*UAA* | Amp | This study |

**Table 3.**
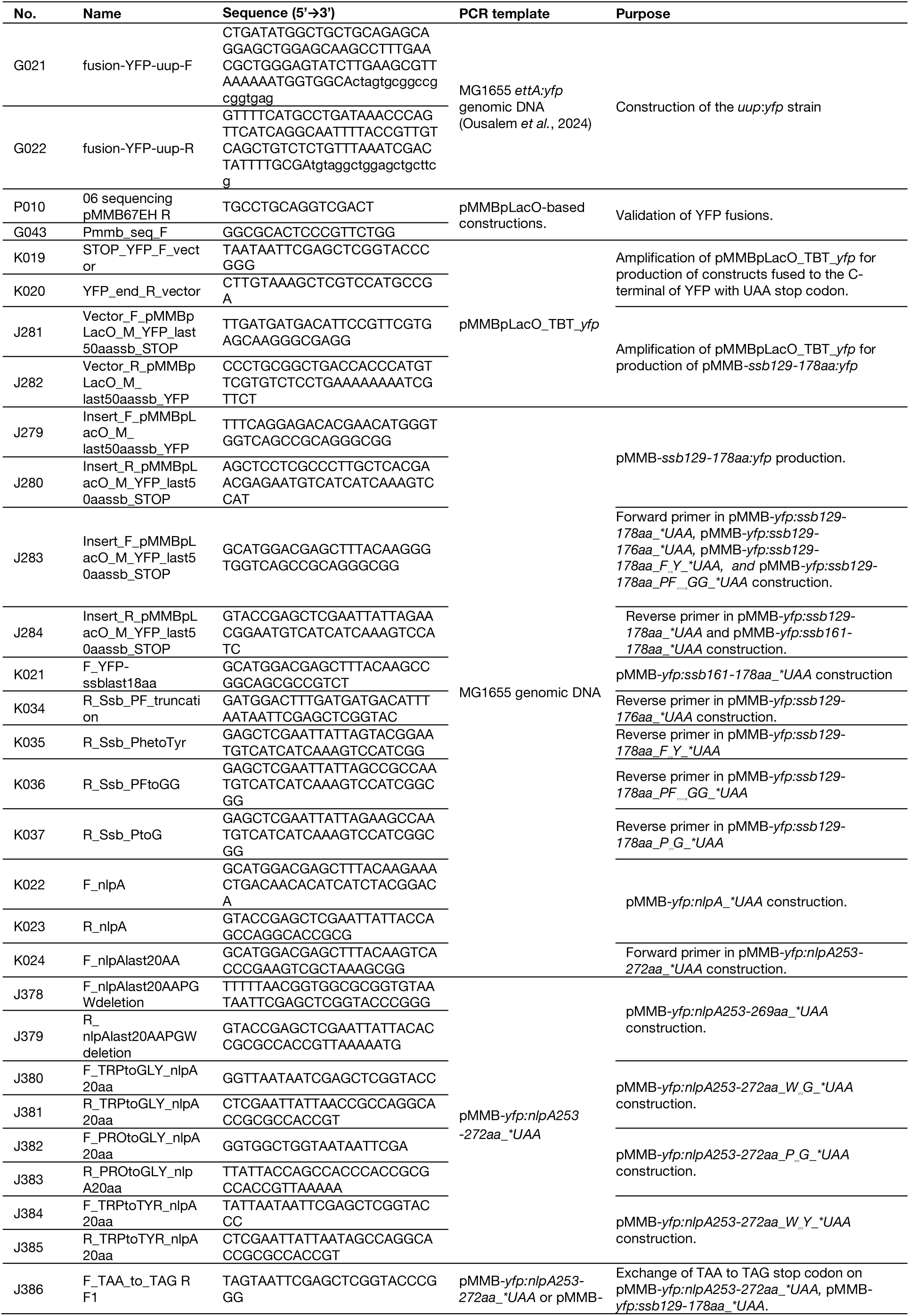
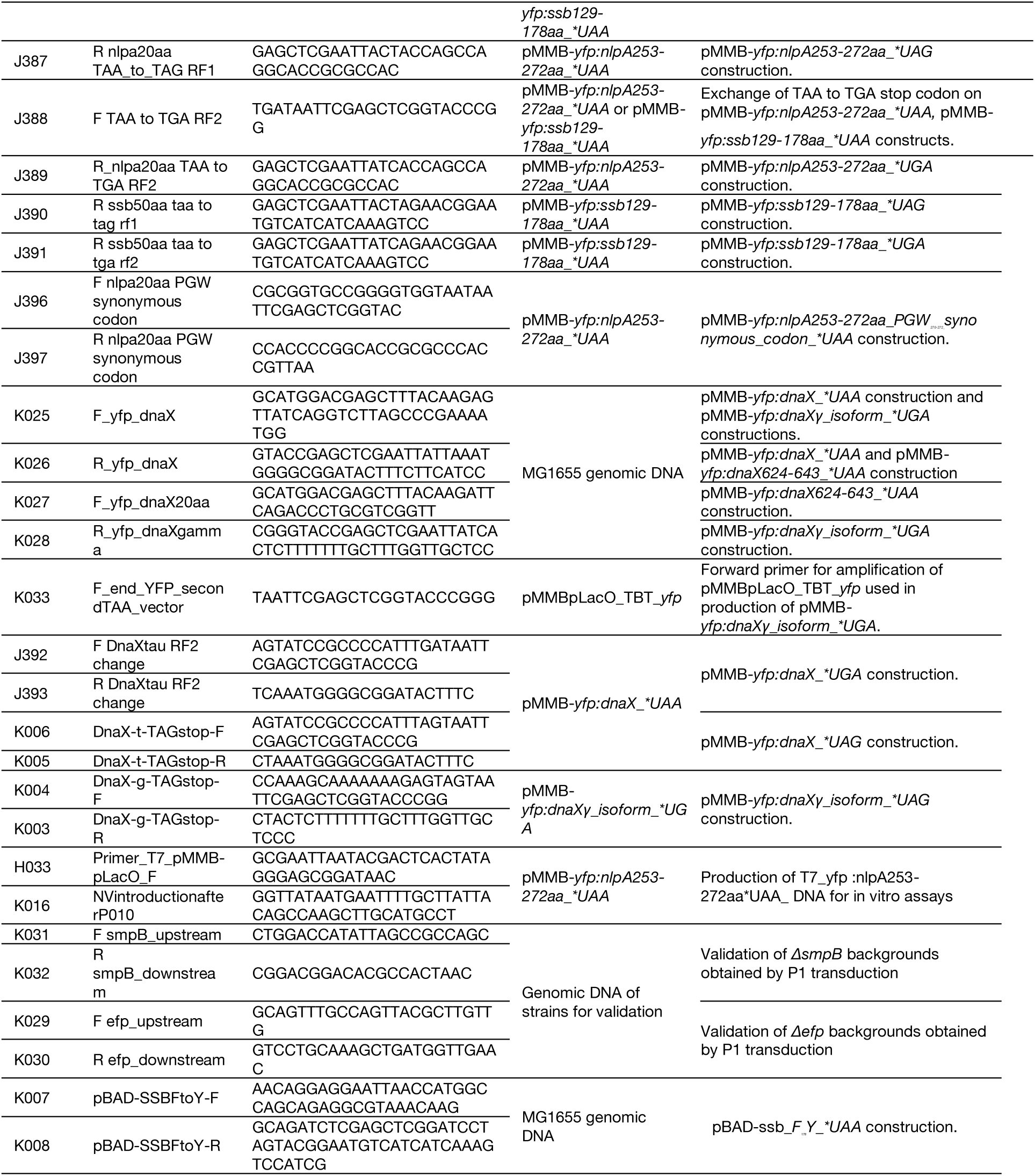
Primers.

## Supporting information

Expended View Figures

## Data availability

RNA-sequencing data are available through the EBI ArrayExpress database (https://www.ebi.ac.uk/biostudies/arrayexpress/studies) under accession code: E-MTAB-17778. Proteomic data are available through the massIVE Repository (https://massive.ucsd.edu/ProteoSAFe/static/massive.jsp) under accession codes: MSV000103466 for the global quantitative proteomic and MSV000103468 for the ssrAH7 tag identification on the YFP:NlpA253-272aa_*UAA product.

## Author contributions

AW with the help of SN and LM performed the experiments. AW, LM and GB analyzed the data. AW, LM and GB prepared the figures. GB designed the research program. GB, AW, LM, wrote the manuscript in consultation with the other authors.

## Acknowledgements

This project has received funding from the European Union’s Horizon 2020 research and innovation program under the Marie Skłodowska-Curie grant agreement No 101034407 through a fellowship to AW. SN, LM and GB were supported by funds from the CNRS (EGM), Université Paris Cité, the LABEX program (DYNAMO ANR-11-LABX-0011) and two ANR grants (EZOtrad/ANR-14-ACHN-0027 and ABC-F_AB/ANR-18-CE35-0010). The proteomic platform was supported by the grant « Equipex Cacsice » (ANR-11-EQPX-0008). We express our great gratitude to Dr. Philippe Bouloc and Hirotada Mori for their gifts of the *ssrAH7* strain and the Keio collection, respectively. We thank Daniel Goicochea Paredes for the construction of some of the YFP reporters and Morgane Wuckelt for the construction of the *uup:yfp* strain.

