## Supplementary material for "The ABC-F protein Uup, Rescues Stalled Ribosomal Termination Complexes to Modulate DNA Maintenance": Expended View Figures

### Expanded View

### For

**Contents:**

Expanded View Figure 1 to 5

### Expanded View Figure 1

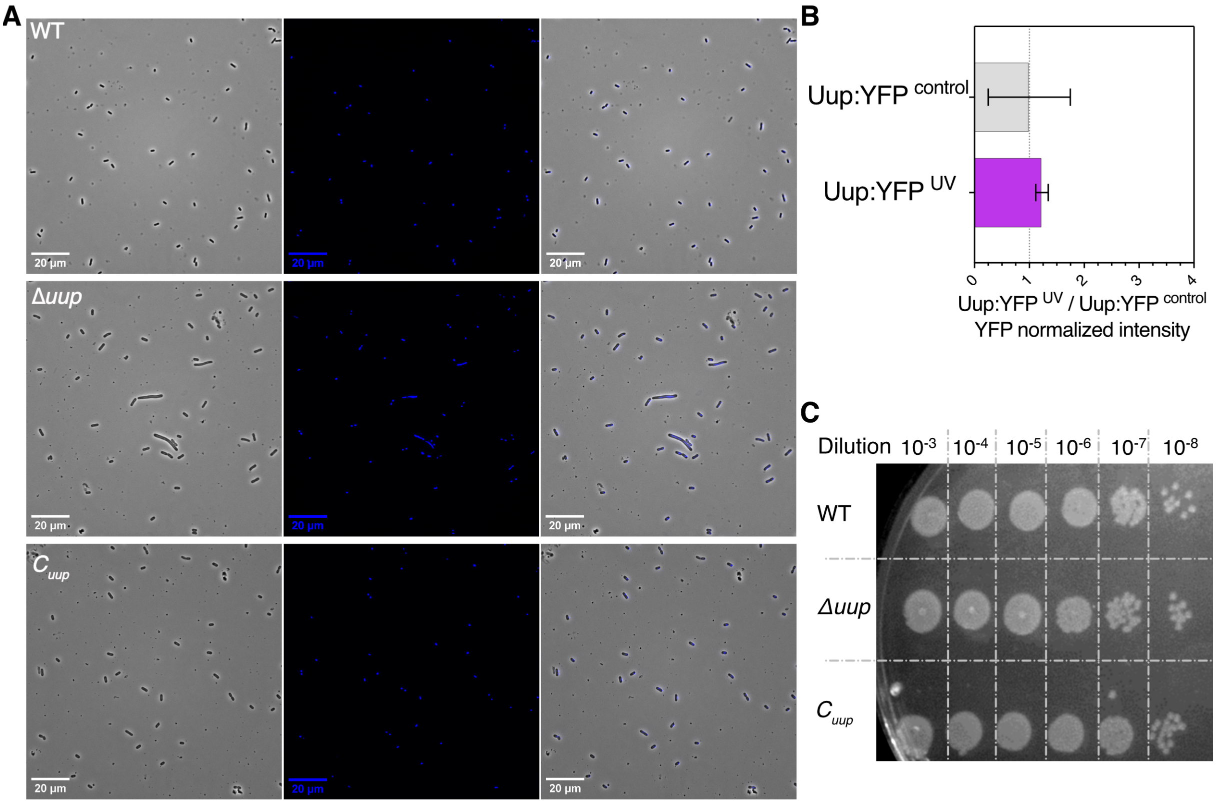


**EV Figure 1: Deletion of *uup* induces phenotypes related to genome instability. A)** Phase-contrast microscopy (left) with DAPI staining (center) and overlay (right) of the WT and Δ*uup* strains from LB culture at $OD_{600}$ = 0.7. **B**) Fluorescence of the *uup:yfp* genomic fusion in cultures at $OD_{600}$ = 1, before UV exposure. Histograms show fluorescence normalized to non-exposed cells ($Uup:YFP^{control}$) before exposure. Error bars represent the mean ± s.d. for triplicate experiments. **D**) Drop dilution assay of WT, Δ*uup*, and $C_{uup}$ strains before UV exposure.

#### Expanded View Figure 2

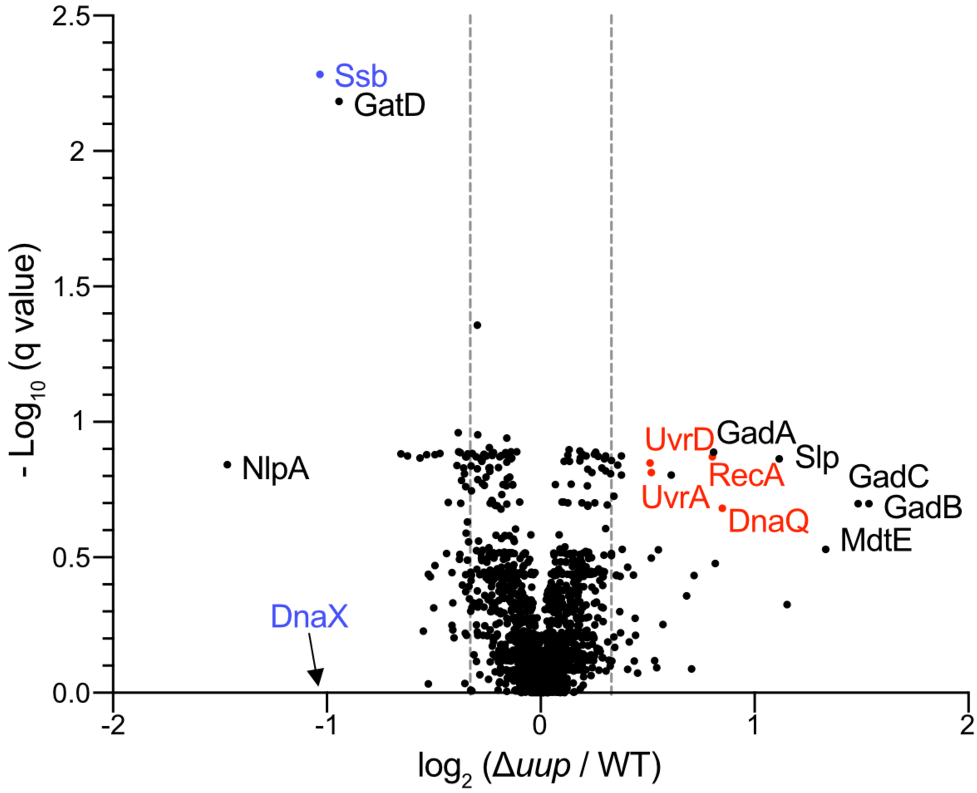


**EV Figure 2: Proteomic analysis of the Δ*uup* strain.** The volcano plot depicts the protein expression profile in the Δ*uup* strain compared to the wild-type (WT) strain grown to an $OD_{600}$ of 2. The ratio of protein quantities (Δ*uup*/WT) and their corresponding *q*-values (corrected *p*-values for false discovery rate using the Benjamini–Hochberg test) are presented on $\log_2$ and $-\log_{10}$ scales, respectively. Proteins differentially expressed in the Δ*uup* strain and involved in the SOS response or DNA maintenance are highlighted in red and blue, respectively. The protein DnaX was not identified in all replicates (injection and biological), and thus no *q*-value is available.

#### Expanded View Figure 3

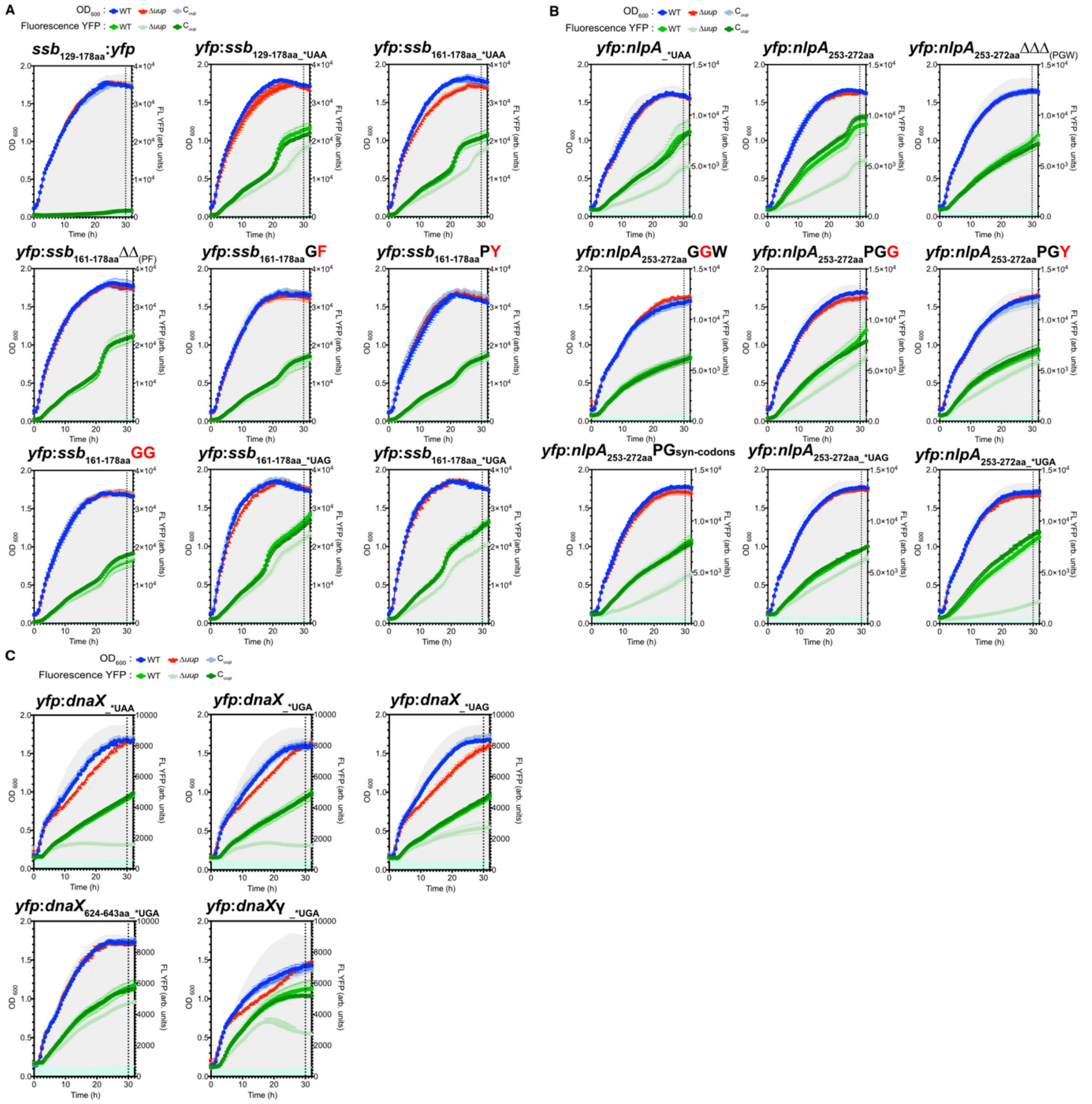


**EV Figure 3: Growth and fluorescence emission curves of *E. coli* strains expressing YFP reporters (Full view of the data presented in figure 4).** Growth curves (blue, cyan, and red) and fluorescence emission curves (green gradient) of triplicate cultures of the WT, Δ*uup*, and $C_{uup}$ strains expressing various YFP reporters. Growth was performed in LB medium supplemented with ampicillin and IPTG to induce reporter expression, except for the non-induced control condition, where no IPTG was added. $OD_{600}$ and fluorescence measurements were recorded every 30 minutes using a ClarioStar microplate reader. The gray and green clouds represent the $OD_{600}$ and fluorescence measurements, respectively, of the WT control strain harboring the plasmid with the YFP reporter under non-induced conditions (without IPTG). Error bars indicate the mean ± s.d. for triplicate experiments. The dotted line at 30 hours corresponds to the fluorescence values reported in **Fig. 4**. **A)** YFP reporters containing parts of the *ssb* coding sequence: the last 50 codons (*ssb*$_{129\text{-}178aa}$) positioned before or after the *yfp* (*ssb*$_{129\text{-}178aa}$:*yfp* and *yfp*:*ssb*$_{129\text{-}178aa}$), the last 18 codons (*ssb*$_{161\text{-}178aa}$) positioned after the *yfp* (*yfp*:*ssb*$_{161\text{-}178aa}$), and variants of the 50-codon construct positioned after the *yfp* (*yfp*: *ssb*$_{129\text{-}178aa}$), where the last amino acids are deleted (ΔΔ)

#### Expanded View Figure 3

or mutated (**G**F, P**Y** and **GG**, mutated residues in bold), and the stop codon is replaced by one exclusively recognized by RF1 (*UAG) or RF2 (*UGA). **B**) YFP reporters containing the full *nlpA* sequence (*yfp*:*nlpA*) or the last 20-codon (*nlpA*$_{253\text{-}272aa}$) positioned after the *yfp* (*yfp*:*nlpA*$_{253\text{-}272aa}$), where the last amino acids are deleted (ΔΔΔ) or mutated (**G**GW, PG**G**, PG**Y**) or replaced by synonymous codon (*yfp*:*nlpA*$_{253\text{-}272aa}$PGsyn-codons), and the stop codon is replaced by one exclusively recognized by RF1 (*UAG) or RF2 (*UGA). **C**) *yfp* reporters containing the full *dnaX* sequence encoding the $\tau$ and $\gamma$ subunits the DNA polymerase III (*yfp*:*dnaX*). YFP reporters with the stop codon replaced by one exclusively recognized by RF1 (*UAG) or RF2 (*UGA) were tested. Variants containing only the last 20 codons of Pol III$\tau$ (*dnaX*$_{624\text{-}643aa}$) were also tested. Finally, a construction where only the $\gamma$ subunit is produced was made (*yfp*:*dnaX*$\gamma$).

#### Expanded View Figure 4

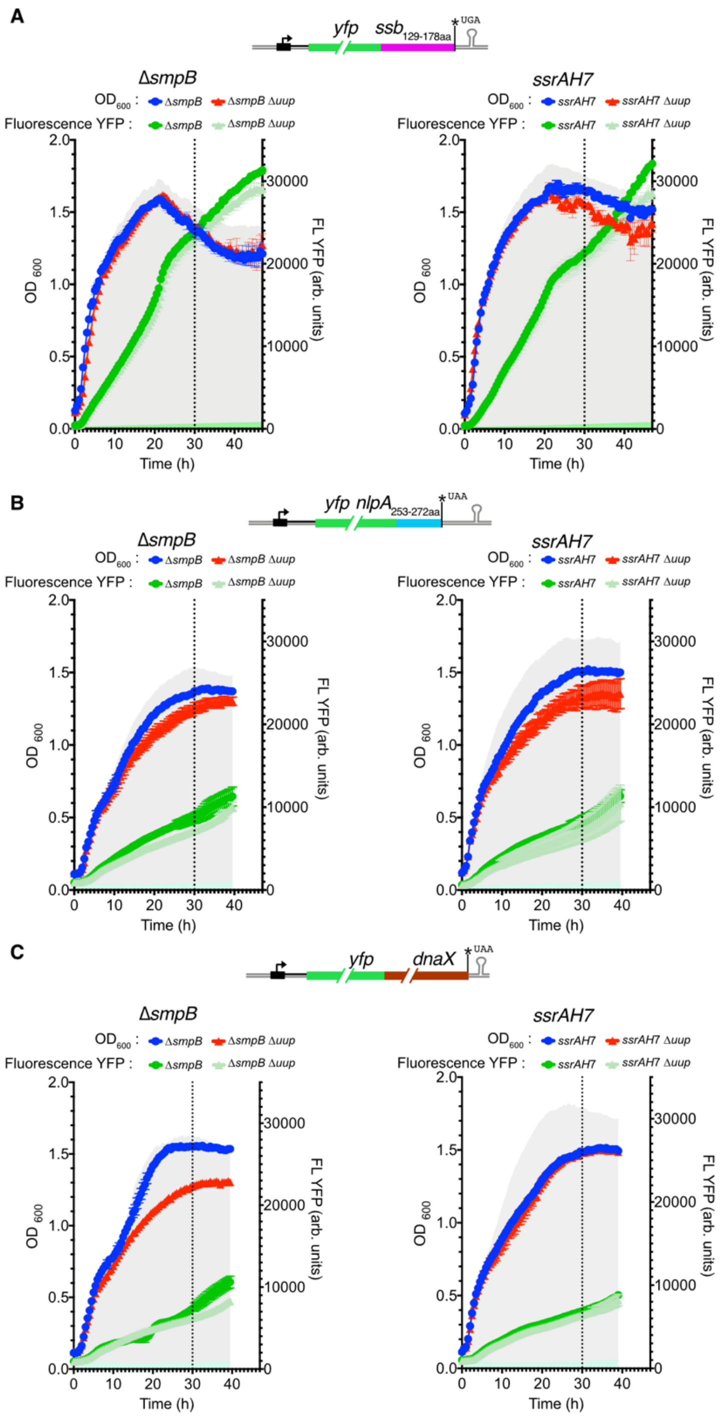


**EV Figure 4: Growth and fluorescence emission curves of *E. coli* strains expressing YFP reporters in different genetic background (Full view of the data presented in figure 6).** Growth curves (blue and red) and fluorescence emission curves (green gradient) of triplicate cultures of Δ*smpB* (left) and *ssrAH7* (right) strains, with or without deletion of *uup* expressing various YFP reporters. Growth was performed in LB medium supplemented with ampicillin and IPTG to induce reporter expression, except for the non-induced control condition, where no IPTG was added. $OD_{600}$ and fluorescence measurements were recorded every 30 minutes using a ClarioStar microplate reader. The gray and green clouds represent the $OD_{600}$ and fluorescence measurements, respectively, of the control strain harboring the plasmid

**Expanded View Figure 4**

with the YFP reporter under non-induced conditions (without IPTG). Error bars indicate the mean ± s.d. for triplicate experiments. The dotted line at 30 hours corresponds to the fluorescence values reported in **Fig. 6**. **A)** Expression of the *yfp*:*ssb*$_{129–178aa}$ reporter. **B)** Expression of the *yfp*:*nlpA*$_{253-272aa}$ reporter. **C)** Expression of the *yfp*:*dnaX* reporter containing the full *dnaX* sequence encoding for both the $\tau$ and $\gamma$ subunits.

#### Expanded View Figure 5

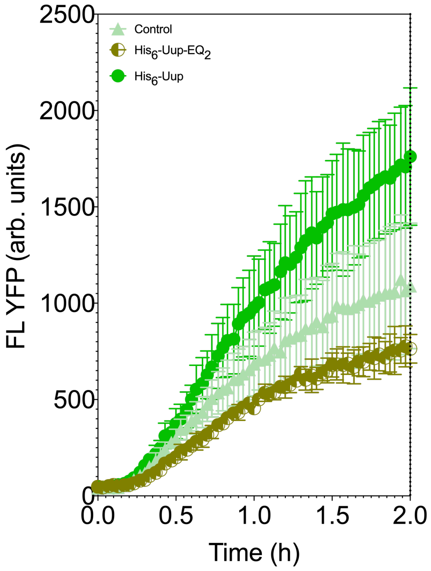


**EV Figure 5: IVTA of the *yfp*:*nlpA*$_{252\text{-}272aa}$ mRNA with or without addition of purified His$_6$-Uup-EQ$_2$ or His$_6$-Uup.** Minimum *in vitro* translation assay using PUREFrex 1.0 cell free protein synthesis kit (GeneFrontier Corporation) in presence of 0.5 µM of purified MRE600 ribosomes and 0.5 µM of the purified *yfp*:*nlpA*$_{252\text{-}272aa}$ mRNA (obtain by T7 *in vitro* transcription). The assay includes the addition of the protein purification buffer (Control), 1 µM of purified His$_6$-Uup-EQ$_2$ or 1 µM of purified His$_6$-Uup. Reactions were performed in triplicate at 37 °C in a microplate reader. Error bars indicate the mean ± s.d. for triplicate experiments.
